# A target expression threshold dictates invader defense and autoimmunity by CRISPR-Cas13

**DOI:** 10.1101/2021.11.23.469693

**Authors:** Elena Vialetto, Yanying Yu, Scott P. Collins, Katharina G. Wandera, Lars Barquist, Chase L. Beisel

## Abstract

Immune systems must recognize and clear foreign invaders without eliciting autoimmunity. CRISPR-Cas immune systems in prokaryotes manage this task by following two criteria: extensive guide:target complementarity and a defined target-flanking motif. Here we report an additional requirement for RNA-targeting CRISPR-Cas13 systems: expression of the target transcript exceeding a threshold. This finding is based on targeting endogenous non-essential transcripts, which rarely elicited dormancy through collateral RNA degradation. Instead, eliciting dormancy required over-expressing targeted transcripts above a threshold. A genome-wide screen confirmed target expression levels as the principal determinant of cytotoxic autoimmunity and revealed that the threshold shifts with the guide:target pair. This expression threshold ensured defense against a lytic bacteriophage yet allowed tolerance of a targeted beneficial gene expressed from an invading plasmid. These findings establish target expression levels as a third criterion for immune activation by RNA-targeting CRISPR-Cas systems, buffering against autoimmunity and distinguishing pathogenic and benign invaders.

**HIGHLIGHTS:**

- Cas13-induced dormancy requires RNA target levels to exceed an expression threshold
- The expression threshold can prevent cytotoxic self-targeting for endogenous transcripts
- The threshold shifts depending on the CRISPR RNA guide:target pair
- The threshold allows cells to distinguish pathogenic and benign infections

## INTRODUCTION

All cellular immune systems face the challenge of differentiating foreign entities from the host. Failure to recognize features associated with an invader exposes the host to a potentially fatal infection, while failure to ignore similar features in the host can trigger a catastrophic autoimmune response. Elucidating how immune systems make these decisions and the consequences of errors is critical not only for treating infectious and autoimmune diseases in higher eukaryotes but also for understanding the physiology and evolution of single-cell microbes under constant assault by mobile genetic elements and bacteriophages (Goldberg and Marraffini, 2015; Hampton et al., 2020; Theofilopoulos et al., 2017).

CRISPR-Cas systems, as the only known adaptive immune systems in bacteria and archaea, face these same challenges (Barrangou et al., 2007; Makarova et al., 2015, 2020). These systems store fragments of invading genetic material as spacers situated between conserved repeats in CRISPR arrays (Barrangou et al., 2007; Jackson et al., 2017). To stem an infection, the arrays are transcribed and processed into individual CRISPR RNAs (crRNAs) that direct the system’s effector nucleases to complementary genetic sequences (Charpentier et al., 2015; Hille et al., 2018; Marraffini and Sontheimer, 2010; van der Oost et al., 2014). The nature of the target genetic material (i.e., DNA or RNA) and the consequence of recognizing a target sequence vary widely across systems. For example, Type II CRISPR-Cas systems recognize target sequences within double-stranded DNA flanked by a protospacer-adjacent motif (PAM) (Leenay and Beisel, 2017), which triggers the Cas9 effector nuclease to create a clean DNA cut (Gasiunas et al., 2012; Jinek et al., 2012). Separately, Type VI CRISPR-Cas systems recognize target sequences within RNA lacking complementarity between the flanking sequence and repeat-derived “tag” within the crRNA (also called a protospacer-flanking sequence or PFS) (Meeske and Marraffini, 2018; Wang et al., 2021). Target recognition by the Type VI Cas13 effector nuclease activates non-specific cleavage of cellular RNAs (Abudayyeh et al., 2016; East-Seletsky et al., 2016). Widespread RNA degradation induces growth arrest--a state called cellular dormancy--that prevents the replication and dissemination of the invader (Abudayyeh et al., 2016; Meeske et al., 2019).

Whether for RNA-targeting or DNA-targeting CRISPR-Cas systems, immune defense is activated in the presence of a nucleic acid complementary to the crRNA guide and flanked by an appropriate sequence. The requirement for target complementarity allows the nuclease to ignore similar but not identical sequences potentially present in the cell. The requirement for the flanking sequence helps the nuclease distinguish the target sequence of the invader from the corresponding spacer within the CRISPR array, as they contain the same complementary sequence but different flanking sequences. However, in the infrequent instance of a spacer being acquired from a chromosomal sequence, this pairing would be expected to drive cytotoxic autoimmunity, killing the cell or driving either mutation of the target or inactivation of the CRISPR-Cas system (Rollie et al., 2020; Stern et al., 2010).

Here, we report that Type VI CRISPR-Cas systems encoding Cas13 nucleases take into account a third criterion for immune activation: minimal expression of the target transcript. This expression threshold is higher than most cellular transcripts under our experimental conditions, allowing the co-existence of a targeted endogenous transcript and an active immune system. The threshold allows the immune system to tolerate a passive invader such as a plasmid yet mount a robust immune response to an actively replicating invader such as a lytic phage. These insights support a model for the impact of Cas13 activation on the cell, where transcripts below the threshold are tolerated or lead to specific target cleavage and transcripts above the threshold lead to widespread RNA degradation. This model may help explain why applying Cas13 in eukaryotes yields programmable gene silencing in some contexts (Abudayyeh et al., 2017; Cox et al., 2017) and non-specific RNA degradation in others (Buchman et al., 2020; Konermann et al., 2018; Wang et al., 2019; Xu et al., 2021).

## RESULTS

### CRISPR-Cas13 fails to induce dormancy when targeting selected endogenous transcripts

Cas13-induced immunity has been principally assessed through targeting transcripts expressed from plasmids or phages (Abudayyeh et al., 2016, 2017; Kiga et al., 2020; Meeske and Marraffini, 2018; Meeske et al., 2019). What has remained less clear is the impact of targeting chromosomally-encoded transcripts, particularly given that targeting chromosomally-encoded transcripts in eukaryotes has been largely associated with targeted gene silencing (Abudayyeh et al., 2017; Cox et al., 2017). To address this gap, we employed the Cas13 nuclease from the Type VI-A CRISPR-Cas system in *Leptotrichia shahii* (LshCas13a) (Abudayyeh et al., 2016; Liu et al., 2017a; Watanabe et al., 2019) in *Escherichia coli* as a simple model of immune defense, paralleling prior studies showing that this nuclease could drive collateral RNA cleavage and dormancy (Abudayyeh et al., 2016; East-Seletsky et al., 2016; Liu et al., 2017b). We first verified Cas13a activity by targeting either of two different sites within a synthetic transcript constitutively expressed from a plasmid, which resulted in a 2,800-fold and 460-fold reduction in the transformation efficiency of the respective crRNA-encoding plasmid versus a non-targeting crRNA plasmid (**Fig. 1A-B**). To verify that the reduction in transformation depended on Cas13a endonuclease activity, we inactivated the HEPN domain of the nuclease using a previously characterized point mutation (R1278A) that abolishes RNA degradation (Abudayyeh et al., 2016). The resulting catalytically-dead Cas13a (dCas13a) yielded similar colony counts between targeting and non-targeting crRNAs, demonstrating that Cas13a RNase activity is responsible for the observed reduction in plasmid transformation (**Fig. 1B**). We further confirmed that targeting the synthetic transcript induces collateral activity based on the degradation of total RNA upon target expression (**Fig. S1**).

**Figure 1.**
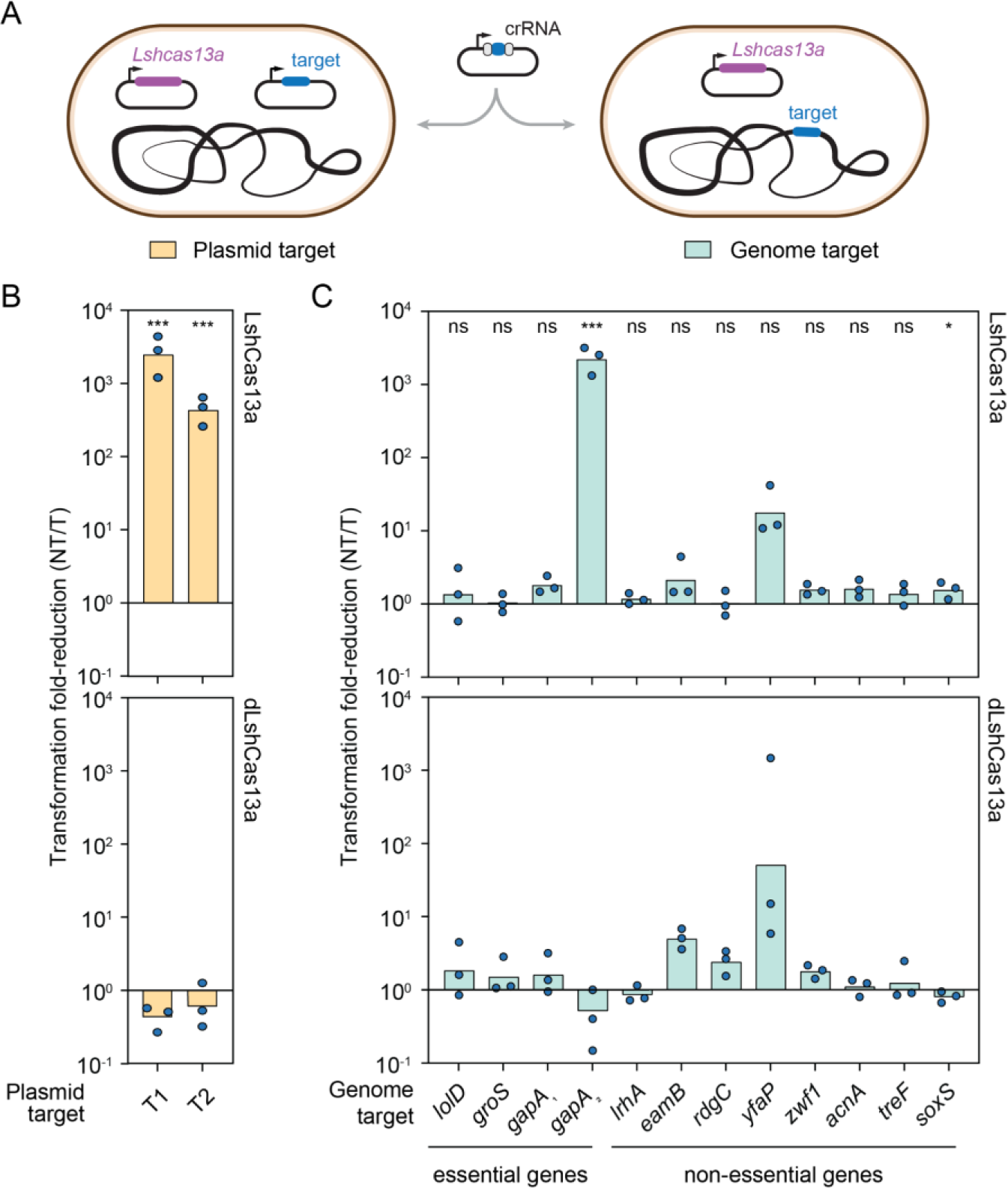
CRISPR-Cas13a is functional in *E. coli* yet fails to confer autoimmunity against selected endogenous targets. (**A**) Experimental setup for the plasmid transformation assay. A crRNA plasmid targeting either a plasmid-encoded or genomically-encoded transcript is transformed into *E. coli* cells expressing the *L. shahii* Cas13a (LshCas13a). The relative number of transformants compared to transformation of a non-targeting crRNA plasmid are quantified. (**B**) Impact of targeting a plasmid-encoded transcript with LshCas13a. dCas13a: catalytically-dead Cas13a. Two different regions were targeted (T1, T2) within a transcript encoding a portion of the *mRFP1* gene. The transcript was expressed from the strong constitutive promoter P8. Fold-reduction was calculated as the ratio of transformants with the non-targeting (NT) and targeting (T) crRNA plasmids. (**C**) Impact of targeting different genomically-encoded transcripts with LshCas13a. Transcripts were targeted from genes considered essential or non-essential in *E. coli* MG1655 under standard growth conditions. *gapA_1_* and *gapA_2_* represent two different target locations within the *gapA* transcript. Bars represent the mean of triplicate independent experiments. Statistical significance was calculated by comparing the transformation fold-reduction for LshCas13a and dLshCas13a. ***: p < 0.001. **: p < 0.01. *: p < 0.05. ns: not significant.

After validating the targeting activity of LshCas13a in *E. coli*, we proceeded to assess self-targeting. Previous studies of Cas13a in bacteria suggested that targeting leads to dormancy as long as the target transcript is present (Abudayyeh et al., 2016; Meeske and Marraffini, 2018), so we expected to observe autoimmunity for all the targets expressed under standard growth conditions. To assess self-targeting, we designed 12 crRNAs targeting mRNAs encoded by essential and non-essential genes in *E. coli* following current guide design rules (Wessels et al., 2020) and repeated the transformation assay (**Fig. 1A**). Surprisingly, targeting yielded a negligible reduction in colony counts for all but two targets: one in the essential *gapA* mRNA and another in the non-essential *yfaP* mRNA (**Fig. 1C**). Repeating the assay with dCas13a or without the nuclease revealed that the transformation reduction with the *yfaP*-targeting crRNA was due to cytotoxicity associated with the crRNA itself (**Figs. 1C** **and S2**). While some Cas13a crRNAs are known to exhibit poor targeting activity (Wessels et al., 2020), the chance that we had selected almost entirely “bad” guides seemed low. Therefore, other factors may be necessary to explain the consistent lack of reduced transformation.

### Boosting target expression above a threshold enables Cas13-based immunity

Given that transcript levels are generally lower when expressed from the chromosome versus a multicopy plasmid, we asked if levels of the target transcript play a role in the induction of Cas13 immunity. We constitutively expressed a 72-nucleotide (nt) fragment of selected mRNA targets on a plasmid (**Fig. 2A**). Expressing these fragments without disrupting the endogenous locus resulted in a 280-fold to 1,000-fold reduction in colony counts compared to the non-targeting control, indicating that target expression levels impact the immune response. The large reduction in colony counts was lost when using dCas13a (**Fig. 2A**), confirming the involvement of RNA cleavage.

**Figure 2.**
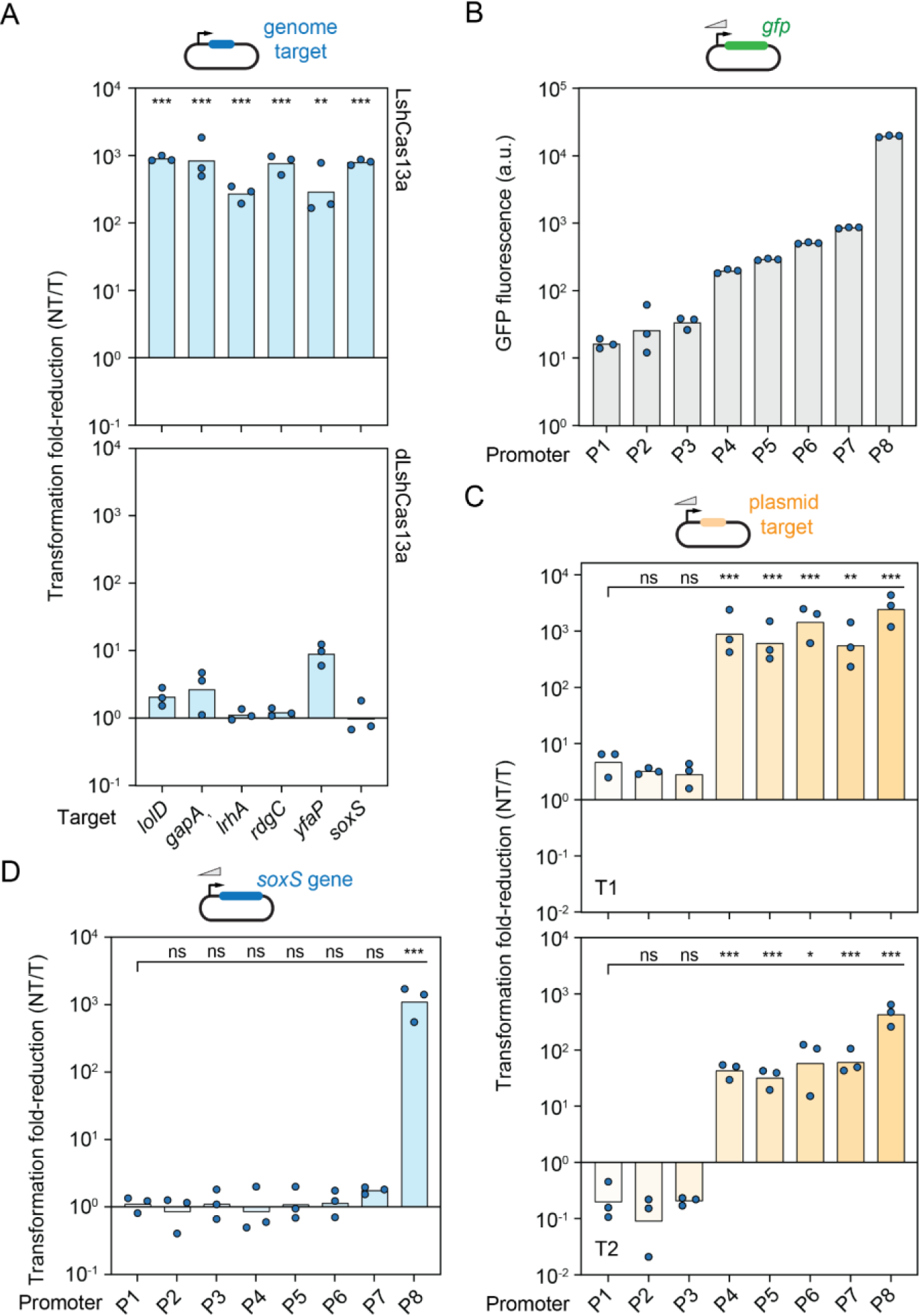
Cas13a-induced immunity requires target expression to exceed a threshold. (**A**) Impact of over-expressing genomic targets in *E. coli*. Targets include the sequence complementary to the crRNA guide along with the 20 nucleotides upstream and downstream and are expressed under the strong constitutive promoter P8. (**B**) Quantified strength of different constitutive promoters in *E. coli*. Promoter strength was measured based on the fluorescence produced from a downstream *gfp* reporter. (**C**) Impact of targeting the plasmid-encoded transcript expressed from different constitutive promoters by Cas13a in *E. coli*. (**D**) Impact of targeting the plasmid-encoded *soxS* transcript expressed from different constitutive promoters by Cas13a in *E. coli*. The genomic copy of *soxS* was intact in the *E. coli* strain. Bars represent the mean of triplicate independent experiments. Statistical significance in A was calculated by comparing the transformation fold-reduction for LshCas13a and dLshCas13a. Statistical significance in C and D was calculated by comparing the transformation fold-reduction to that of the weakest P1 promoter. ***: p < 0.001. **: p < 0.01. *: p < 0.05. ns: not significant.

Our observations suggested that a certain target expression level might need to be reached to elicit the immune response. To explore this possibility, we expressed the plasmid-encoded synthetic transcript targeted at two locations (T1, T2) under eight different constitutive promoters (P1 to P8) (**Table S1**) exhibiting varying expression strengths (**Fig. 2B-C**). While the three weakest promoters yielded no reduction in transformation efficiency compared to the non-targeting control, we observed a strong reduction for the remaining promoters (∼1,100-fold for T1, ∼60-fold for T2). No reduction was observed with dCas13a for any of the promoters (**Fig. S3**). The transition between low and high transformation efficiencies was remarkably sharp and occurred for both targets between the same two promoters separated by only a 5.8-fold difference in transcriptional activity (**Fig. 2C**). We performed a similar analysis with the full-length *soxS* mRNA using the same set of constitutive promoters (**Fig. 2D**), where the transformation reduction occured only with the strongest promoter. These results show that low target expression prevented immune induction, while boosting target expression beyond a threshold was sufficient to induce Cas13-based immunity.

### A genome-wide screen establishes target expression levels as the principal determinant of immune induction

Thus far, our observations were based on a small number of endogenous targets. In order to determine to what extent expression thresholds influence targeting across the transcriptome and whether other factors (e.g., target position, sequence context) also impact the outcome of targeting, we performed a genome-wide screen. We designed a library of 25,597 crRNAs targeting all chromosomally-encoded mRNAs and rRNAs in *E. coli* (**Fig. 3A**). The guide sequences were selected following current design rules (e.g., PFS lacking complementarity to crRNA repeat tag) (Meeske and Marraffini, 2018; Wessels et al., 2020) to reduce the inclusion of low-efficiency guides, and the targets were spaced across the entire length of each coding region (or transcribed region in the case of rRNAs). The library included 400 randomized guides as non-targeting controls lacking complementarity to any endogenous transcripts. We then transformed the crRNA library into *E. coli* cells with or without LshCas13a and cultured the transformed cells with antibiotic selection to deplete guides causing collateral activity and dormancy. Short-read sequencing was finally applied to measure the depletion of each guide compared to the no-LshCash13a control cultured under the same conditions (**Fig. 3B**). Under this setup, highly active guides would be heavily depleted within the library, while poorly active guides would be minimally depleted.

**Figure 3.**
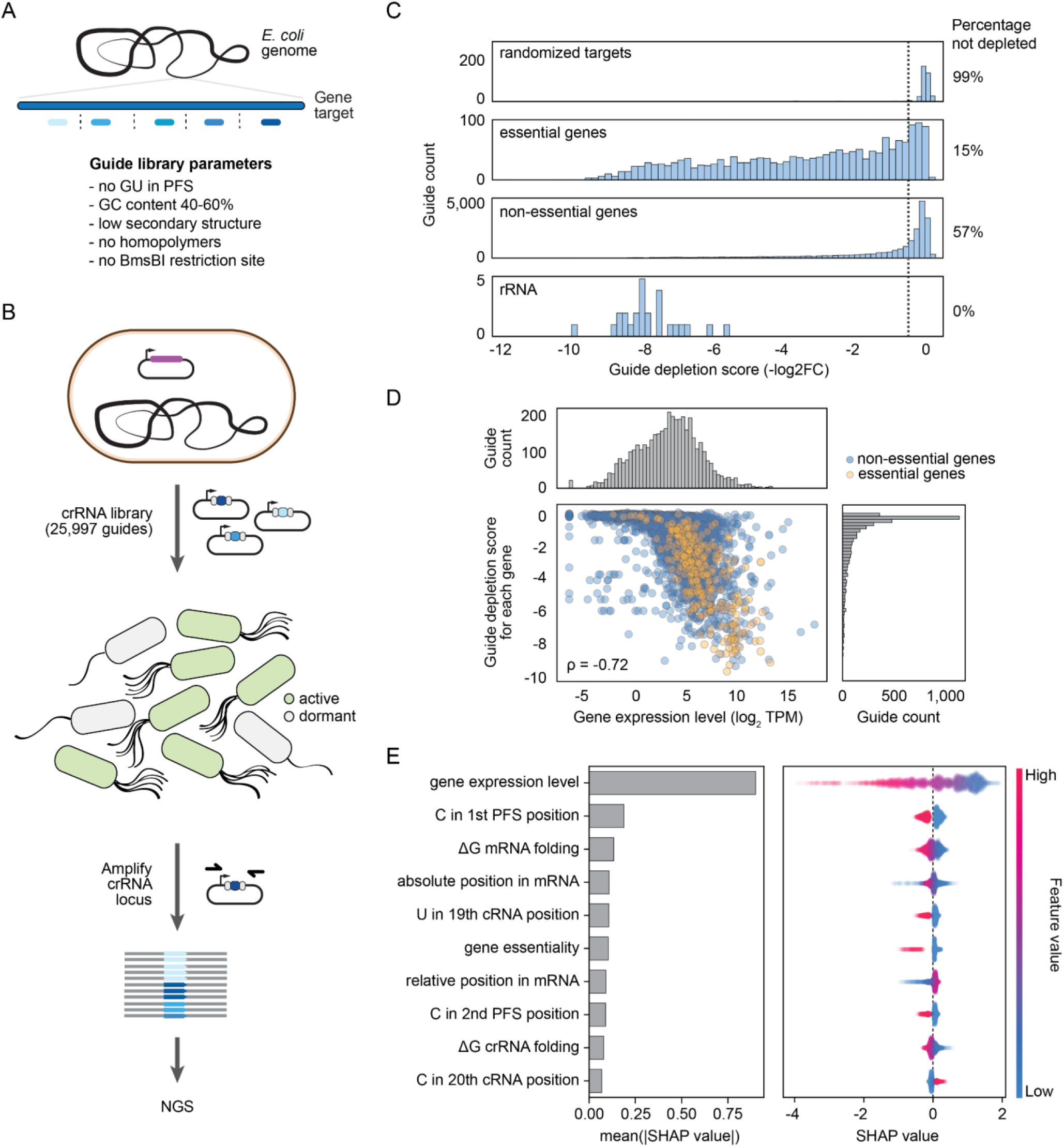
A genome-wide CRISPR-Cas13a screen reveals target expression levels as the main determinant of cytotoxic self-targeting. (**A**) Design of crRNA guide library. Guide selection accounted for standard rules lending to efficient targeting and spanning the entire coding region of each target gene. Other parameters (i.e., homopolymers, BsmBI sites) were included to facilitate library synthesis and cloning. The resulting library included 25,470 guides targeting protein-coding genes, 127 guides targeting rRNAs, and 400 randomized guides as negative controls. (**B**) Workflow for library screening. As part of the screen, cells with or without a LshCas13a plasmid (purple) are transformed with the crRNA plasmid library and cultured while selecting all present plasmids. Guide depletion is determined in comparison to the same workflow with the no-LshCas13a control. (**C**) Distribution of depletion scores for different groups of guides within the library. The cutoff for no fitness defect is based on the range of depletion scores for a set of randomized guides. (**D**) Correlation between guide depletion score and the expression levels of the target gene. Expression levels were measured by RNA-seq analysis with *E. coli* cells harboring the no-LshCas13a control and subjected to the library workflow to a turbidity of ABS_600_ ≈ 0.5. Values for transcript levels and guide depletion are the average of duplicate independent experiments and screens, respectively. ρ: Spearman coefficient. See **Fig. S5A** for the correlation for a turbidity of ABS_600_ ≈ 0.8. (**E**) SHAP values for the strongest predictors of guide depletion from the library. The left barplot indicates the average absolute contribution of each feature to the predicted depletion values, while the right beeswarm plot shows the impact of each feature on each individual prediction.

Initial analysis revealed that the extent of depletion depended on whether the target was associated with an essential or non-essential gene. This observation is in line with the idea that target transcript cleavage of an essential gene would be expected to cause a fitness defect even in the absence of widespread collateral cleavage. Indeed, compared to the set of randomized guides, 85% of guides targeting essential genes were depleted versus only 43% of guides targeting non-essential genes. The median depletion was also higher for guides targeting essential (2.9-fold) versus non-essential (0.3-fold) genes (**Fig. 3C**). The screen was validated by testing individual highly depleted guides using the transformation assay (**Fig. S4**). The limited depletion of guides targeting non-essential genes further shows that guides, even when designed following standard rules, infrequently lead to widespread collateral cleavage and dormancy.

We also noticed that guides targeting rRNAs, the highest expressed RNAs in the cell, were strongly and consistently depleted in the library (**Fig. 3C**). To analyze in more detail how transcript levels contributed to guide depletion across the library, we performed transcriptomics analyses of *E. coli* cells not expressing LshCas13a and cultured in middle (ABS_600_ ≈ 0.5) and late (ABS_600_ ≈ 0.8) exponential growth phases. By correlating the resulting transcript levels with the median depletion of guides targeting each gene, we found a clear correlation that was stronger for transcript levels in middle exponential growth (Spearman coefficient = 0.72) than for late exponential growth (Spearman coefficient = 0.68) (**Figs. 3D** **and S5A**). The stronger correlation with early exponential growth might be because this growth phase dominates the screen. Translational strength predicted using the ribosome-binding site (RBS) calculator showed minimal correlation (Salis, 2011) (**Fig. S5B**), indicating that protection of the mRNA by translating ribosomes does not account for differences in guide depletion.

In order to determine which features of the guide:target pair most impact induction of cytotoxic immunity, we subjected the dataset to machine learning (see Methods) (**Fig. 3E**). The resulting SHAP values (Lundberg et al., 2020) revealed that transcript levels were the strongest predictor of guide depletion. In contrast, gene essentiality had a modest effect on the predicted depletion for the average guide, which can be attributed to the fact that there are only a small number of essential genes in *E. coli*. Factors related to the sequence and predicted structure of the crRNA guide and RNA target also had predictive value (**Figs. 3E** **and S6**), in line with previous observations from high-throughput assays of Cas13a guides (Wessels et al., 2020). Together, our results demonstrate that most self-targeting guides do not activate a Cas13-based immune response, at least under our experimental conditions, but that high transcript levels can trigger immunity.

### The target expression threshold depends on the selected guide

Although transcript levels are an important predictor of immune activation, guide features also impact the response. For example, depletion of the highly expressed *tolB* transcript (**Fig. 4A**), which was targeted by nine different guides within the library, varied between minimal depletion to depletion paralleling that of some guides targeting rRNAs (**Figs. 3C and 4B**). We tested each guide individually in the transformation assay and observed a strong correlation between the reduction in colony counts and the depletion score from the screen (Spearman coefficient = - 0.87). Furthermore, some of the guides yielded an insignificant reduction in colony counts compared to the non-targeting control, underscoring the variability in immune activation for this one highly expressed mRNA depending on the guide sequence.

**Figure 4.**
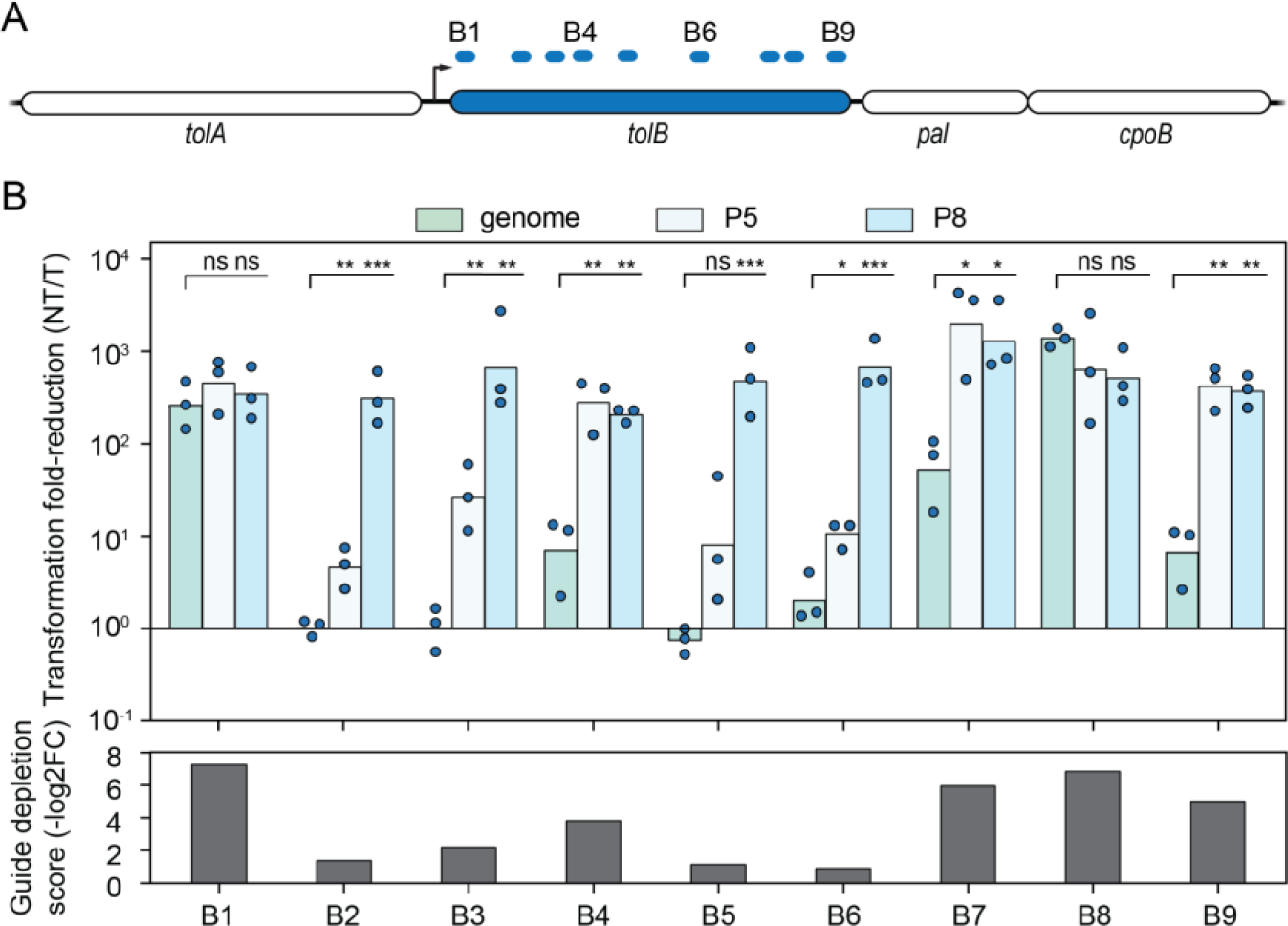
The target expression threshold varies between guide:target pairs associated with the same transcript. (**A**) Target locations within the *tolB* gene from the guide library. Targets are labeled B1 - B9. (**B**) Impact of boosting *tolB* expression for each crRNA guide in *E. coli*. The *tolB* gene was expressed from one of two constitutive promoters (P5 or P8) on a plasmid. The genomic copy of *tolB* was always intact. Bottom: Average guide depletion score for each guide from the library screen. Bars represent the mean of triplicate independent experiments. Statistical significance was calculated by comparing the transformation fold-reduction to that with only the genomic copy of *tolB*. ***: p < 0.001. **: p < 0.01. *: p < 0.05. ns: not significant.

Our initial data also suggested that boosting expression of the *tolB* mRNA might be able to rescue immune activation for poorly performing guides. We therefore placed the *tolB* RBS and coding region under a medium or strong constitutive promoter on a plasmid, and we repeated the transformation assay (**Fig. 4B**). For all guides that displayed low efficiency when targeting the endogenous transcript, expressing *tolB* from the plasmid led to full immune activation. Notably, guides that exhibited some reduction in colony counts required only the medium promoter to maximize immune activation, while guides exhibiting negligible reduction in colony counts required the strong promoter to achieve full immunity. Therefore, boosting target expression can rescue immune activation by otherwise poorly active crRNAs. Concurrently, the target expression threshold required for immune activation by Cas13a can vary between guides, even when targeting the same transcript.

### Reducing Cas13a levels precludes immune induction

We demonstrated that the expression level of the target must exceed a threshold for Cas13a to induce dormancy. While thus far we focused on the properties of the target, the concentration of the Cas13a:crRNA complex could also be an influencing factor. We therefore explored the impact of altering levels of this complex on plasmid transformation. Specifically, we swapped the native promoter from *L. shahii* driving transcription of *cas13a* with a constitutive synthetic promoter with weaker expression (Pw). After replacing *cas13a* with *deGFP* and measuring fluorescence of the cells, we measured a 44-fold decrease in fluorescence for Pw compared to the native promoter (**Fig. S7A**). We then performed the transformation assay targeting the synthetic transcript expressed from the set of constitutive promoters. Remarkably, we did not observe immune activation even when expressing the synthetic target from the strongest promoter (**Fig. S7B**).

To confirm that the nuclease is expressed, we introduced plasmids encoding the nuclease, targeting or non-targeting crRNA and a deGFP reporter to measure collateral cleavage activity in cell-free transcription-translation (TXTL) reactions (Liao et al., 2019; Marshall et al., 2018, 2020) (**Fig. S7C**). Monitoring reporter fluorescence over time, we found that the nuclease expressed from the Pw promoter inhibited deGFP expression compared to the non-targeting control, albeit with slower kinetics than the construct with the native promoter. Nevertheless, these data indicate that Cas13a expressed under the Pw promoter is active. Overall, these results show that low levels of Cas13a nuclease can impair immune activation, even when target expression levels are high.

### Infection by a lytic phage activates Cas13a-based immunity even for less efficient guides

Beyond self-targeting, another important question is how the target expression threshold impacts immune activation by foreign invaders targeted by Cas13. We began with the lytic MS2 RNA bacteriophage that rapidly replicates and lyses *E. coli* as part of the infection cycle. We designed 6 crRNA guides (M1-M6) targeting within the *rep* and *cp* genes (**Fig. 5A**), as both genes are non-toxic when expressed in *E. coli* and thus can be expressed individually to determine the expression threshold for each crRNA. To determine each crRNA’s expression threshold for immune induction, we expressed *rep* or *cp* under a weak, medium, and strong promoter on a low-copy plasmid and measured the extent of plasmid interference (**Fig. 5B**). We found that the crRNAs were associated with a wide range of target expression thresholds for immune induction, with one exhibiting a low expression threshold, three exhibiting a higher expression threshold, and two exhibiting an expression threshold higher than the strongest promoter.

**Figure 5.**
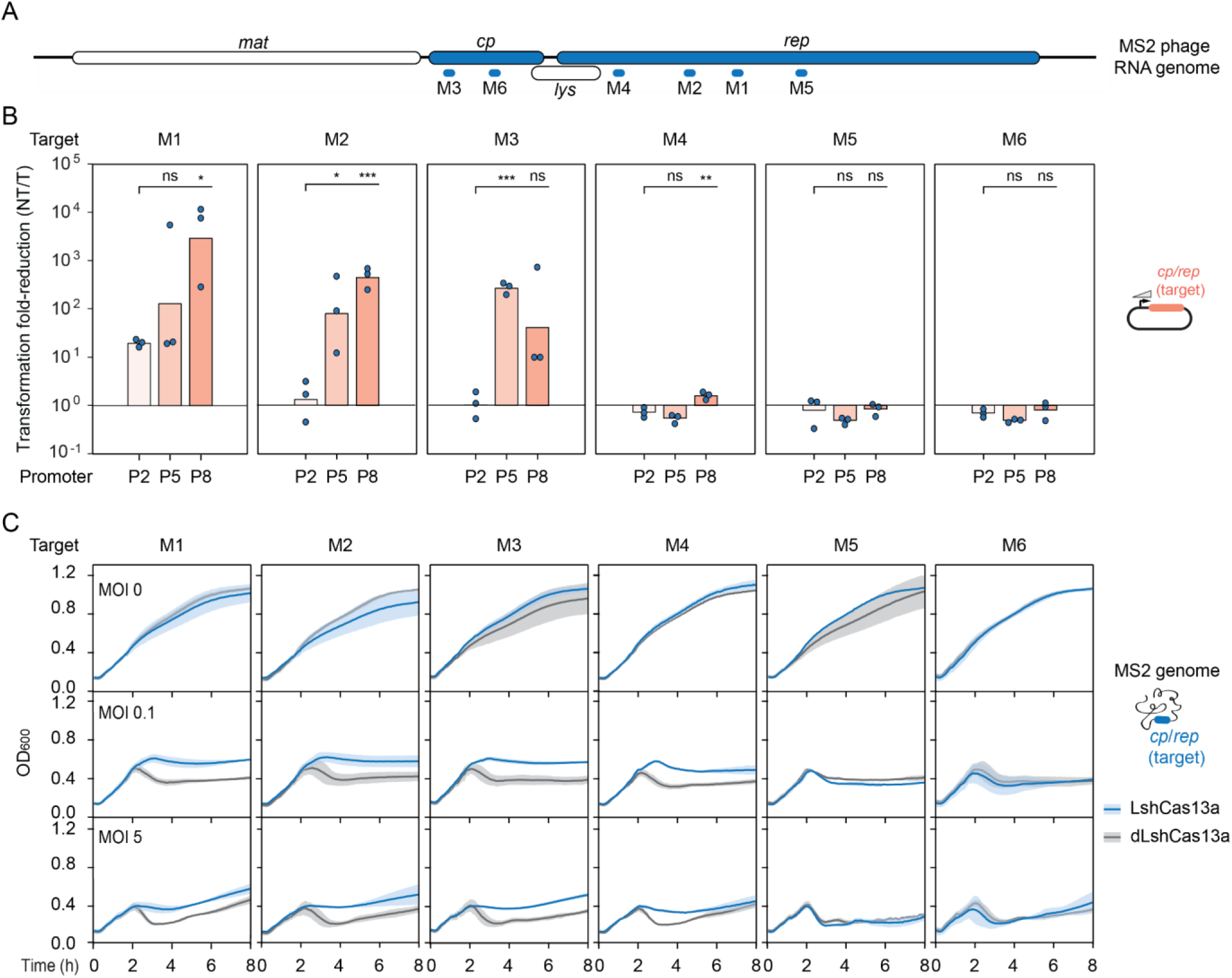
Cas13a defends *E. coli* from a lytic bacteriophage even with guides exhibiting a higher immunity threshold. (**A**) ssRNA genome of the lytic MS2 bacteriophage and location of guides targeting *cp* and *rep* genes used for accessing targeting and phage defense. (**B**) Immunity threshold for different LshCas13a guides targeting *cp* or *rep* genes expressed under different promoters on a plasmid. Bars represent the mean of triplicate experiments. Statistical significance in B was calculated by comparing the transformation fold-reduction to that of the weakest P2 promoter. ***: p < 0.001. **: p < 0.01. *: p < 0.05. ns: not significant or average below the reference. (**C**) LshCas13a defense against MS2 bacteriophage at different MOIs. Growth curves of *E. coli* with active or dead LshCas13a are compared over time. The lines and bars represent respectively the mean and standard deviation of three independent biological replicates.

We then infected *E. coli* cultures with MS2 phages at different MOIs (0.1, 5) and assessed defense through each of the six crRNAs (**Fig. 5C**). In the absence of MS2 phage, all cultures exhibited continual growth over the time course. Cultures with LshCas13a and crRNAs exhibiting at least some immunity in the transformation assay (M1-M4) halted growth and maintained turbidity, indicative of immunity-induced dormancy. In contrast, cultures with LshCas13a and the crRNAs that did not exhibit immunity in the transformation assay (M5, M6) as well as all dLshCas13a controls showed a temporary drop in turbidity, indicative of successful phage infection. Therefore, even crRNAs requiring high target expression to activate widespread immunity can confer robust defense against a pathogenic invader.

### The target expression threshold allows tolerance of invaders conferring benefits to the host

Beyond pathogenic invaders, we considered benign invaders that do not rapidly replicate and instead generally maintain transcript levels. We reasoned that the target transcript would not elicit an immune response if expressed below the expression threshold, allowing the invader to persist. One notable scenario is a targeted gene that benefits the host cell, such as a virulence factor or an antibiotic resistance marker spread through mobile genetic elements (Frost et al., 2005; Koonin et al., 2020; Partridge et al., 2018). If targeting does not substantially affect levels of this transcript, then the host could benefit from its expression but not elicit an immune response.

To explore this scenario directly, we created a plasmid encoding two resistance genes: a kanamycin resistance gene targeted by two distinct crRNAs (our targeted beneficial gene) as well as a non-targeted hygromycin resistance gene (for plasmid selection). The kanamycin gene was placed downstream of a strong, medium, or weak constitutive promoter (**Fig. 6A**). We then assessed tolerance to and growth benefits from the kanamycin resistance gene in two separate steps: assessing tolerance by transforming the plasmid under hygromycin selection, and assessing the growth benefit by measuring growth of tolerant cells in the presence of kanamycin. For the first step, high target expression induced an immune response while low target expression was tolerated (**Fig. 6B**), in line with other target transcripts expressed from plasmids (**Figs. 2, 4**). For the second step, we found that the tolerated crRNA:promoter combinations yielded substantial growth on 10 μg/mL kanamycin (**Fig. 6C**). One combination (K2 crRNA with P5-expressed target) maintained growth even in the presence of 50 μg/mL kanamycin (**Fig. S8**). In contrast, a plasmid lacking the kanamycin resistance gene did not yield any growth. For some combinations, growth on kanamycin was slower with the targeting versus non-targeting crRNA. As this same growth defect was not observed in the presence of hygromycin, the growth defect on kanamycin may be attributed to Cas13a-mediated gene silencing that sensitized the cells to kanamycin. Overall, these results show that the target expression threshold can allow tolerance of a benign invader, even allowing the targeted gene to provide benefits to the host cell.

**Figure 6.**
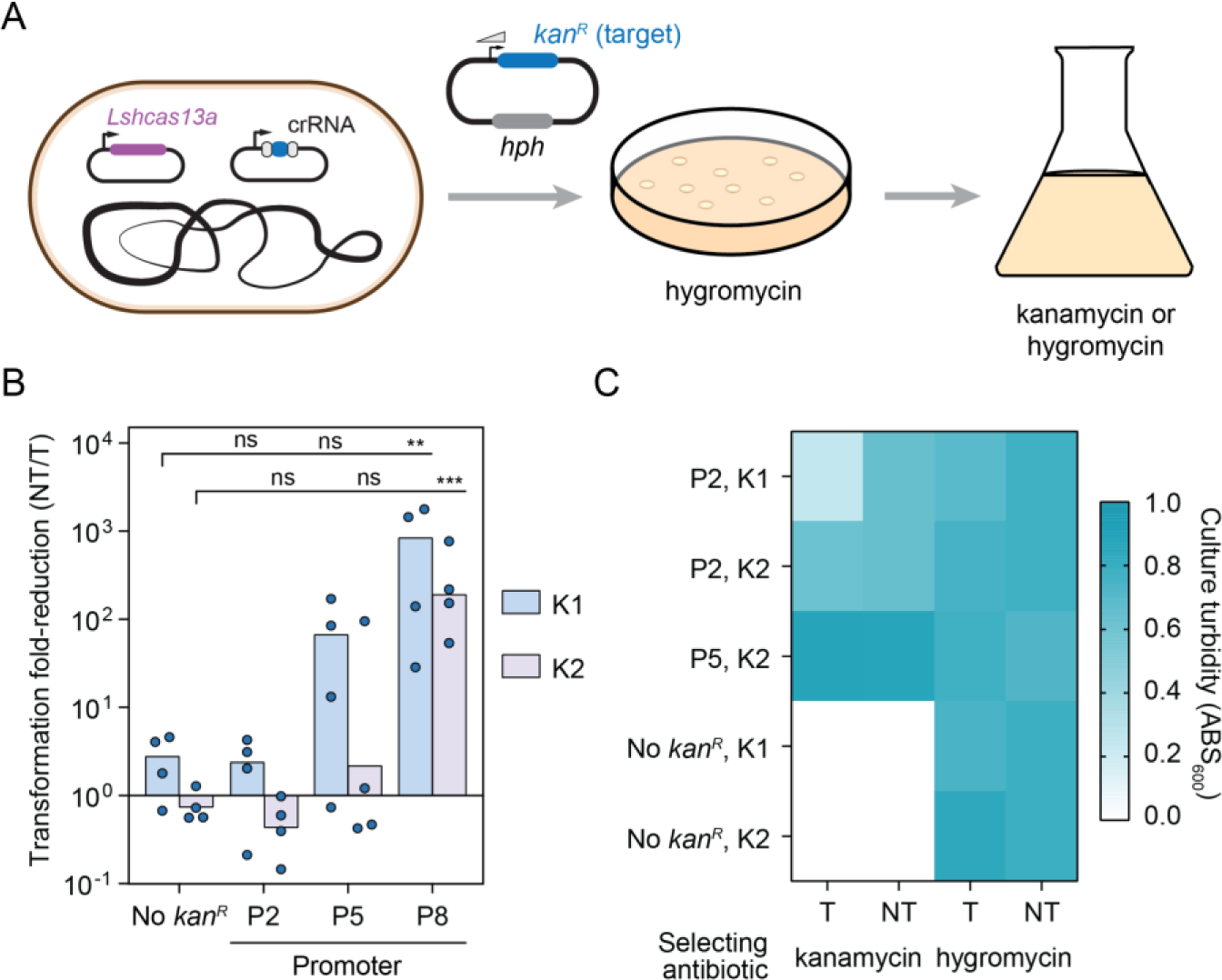
A plasmid transcript targeted by Cas13a and expressed under the threshold can be tolerated and can confer benefits to the host. (**A**) Experimental setup for evaluating tolerance for a plasmid-expressed beneficial gene. The two targets (K1, K2) fall within the *kan^R^* transcript, which is expressed under different constitutive promoters (P2, P5, P8). (**B**) Impact of Cas13a-based targeting of the *kan^R^* transcript expressed under the different constitutive promoters. Bars represent the mean of quadruplicate independent experiments. Statistical significance was calculated by comparing the transformation fold-reduction to that without the *kan^R^* gene. ***: p < 0.001. **: p < 0.01. *: p < 0.05. ns: not significant. (**C**) Heat map comparing growth in the presence of kanamycin (10 μg/mL) or hygromycin (100 μg/mL) for constructs associated with a low transformation fold-reduction in B. Values represent the average turbidity after 12 hours of growth from 9 biological replicates. Constructs lacking the *kan^R^* gene serve as negative controls, while the non-targeting crRNA plasmid serves as a non-targeting (NT) control. See **Fig. S8** for the same growth measurements with a higher concentration of kanamycin (50 μg/mL).

## DISCUSSION

Immune activation by CRISPR nucleases normally requires two established criteria: complementarity between the crRNA guide and the target, and a PAM or PFS flanking the target (Leenay and Beisel, 2017). Here, we show that the expression levels of the target transcript represent a third criterion determining immune activation by the RNA-targeting nuclease Cas13. The outcome depends on whether expression levels are above or below a specific threshold (**Fig. 7**). When target transcript levels are above the threshold, widespread collateral RNA cleavage leads to cell dormancy. When target transcript levels are below the threshold, cells escape dormancy. In that case, target expression can be either unperturbed or silenced. Gene silencing under the expression threshold is supported by two lines of evidence: extensive depletion of guides targeting essential genes but not non-essential genes in the library screen (**Fig. 3C**) and reduced growth on kanamycin but not hygromycin when targeting the *kan^R^* gene (**Figs. 6C** **and S8**). In both cases, silencing of the essential gene would reduce growth, even if the cells do not enter dormancy through collateral RNA cleavage. The target expression threshold also can vary between guide:target pairs, even when the targets are present in the same transcript. With sufficient expression of the target transcript, however, the threshold can be crossed to activate immunity.

**Figure 7.**
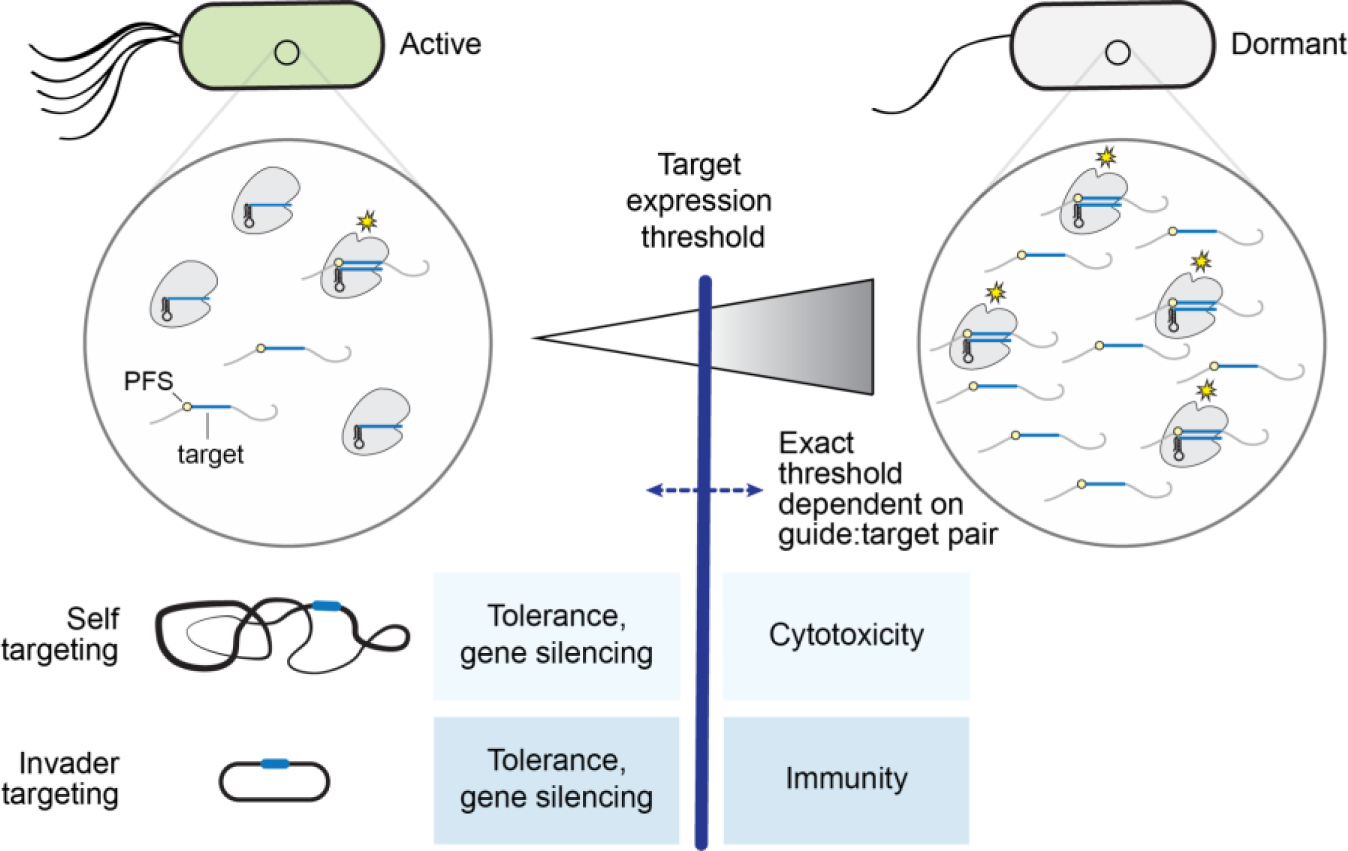
Proposed model for the target expression threshold as a determinant of Cas13-based defense and autoimmunity. Any transcript with extensive complementarity to the crRNA guide flanked by a PFS can be targeted by Cas13. However, the impact of targeting on the target gene and the host cell depends on a third factor: a target expression threshold. Target expression under the threshold is tolerated and can lead to targeted gene silencing, as supported by the extensive depletion of guides targeting essential genes but not non-essential genes in the library screen (Fig. 3) as well as reduced growth on kanamycin while targeting the *kan^R^* resistance gene (Fig. 6C **and S8**). Target expression above the threshold leads to cytotoxicity for self-targeting and immunity for invader targeting. The exact threshold is target-dependent, likely due to factors that influence guide performance (e.g., local secondary structure, GC content).

One simple biochemical explanation for the threshold is the level of activated Cas13:crRNA complexes. Higher levels would lead to widespread RNA cleavage that induces growth arrest and dormancy. Lower levels might lead to preferential cleavage of the target, resulting in specific silencing. For less efficient targets, the target transcript may be poorly recognized for multiple reasons associated with guide selection (e.g., GC content, crRNA or target folding) (Wessels et al., 2020), resulting in a negligible impact on target transcript levels. In some cases, we observed a sharp threshold, where immunity was fully activated with as little as a 5.8-fold change in promoter activity driving target expression. While the extent of collateral cleavage should scale with the concentration of activated Cas13:crRNA complexes, collateral cleavage could create a positive feedback loop. Specifically, degradation of key transcripts might reduce growth, leading to slowed dilution of activated Cas13:crRNA complexes (Elowitz et al., 2002), leading to further RNA degradation and growth arrest. This mechanism is supported by our findings that lowering Cas13a levels prevents immune induction (**Fig. S7**), where the concentration of activated Cas13a:crRNA complexes could not be achieved to initiate the feedback loop. However, there could also be other possible explanations for the sharp threshold, such as cooperativity between Cas13 enzymes or how well activated complexes can diffuse through the dense cytoplasm of a bacterial cell. Future work could evaluate the link between collateral RNA cleavage and growth arrest as well as other factors that affect the threshold, such as expression levels of the crRNAs, the impact of multiplexed targeting, and the extent of collateral activity exhibited by different Cas13 nucleases (Abudayyeh et al., 2017; Xu et al., 2021).

The existence of a target expression threshold has major implications for self-targeting and invader defense (**Fig. 7**). For targets above the threshold, immunity resembles that associated with other CRISPR-Cas systems: cytotoxicity incompatible with cell survival when targeting the host’s own genetic material, or invader clearance or induced dormancy when targeting an invader. For targets below the threshold, immunity takes a different form. For self-targeting, targets under the threshold are tolerated, allowing the cells to persist with a limited impact on fitness. In the event the target transcript undergoes silencing, Cas13 could serve a regulatory role without compromising its ability to conduct immune defense, similar to scaRNAs and Cas9 (Ratner et al., 2019). For invader defense, the threshold ties immune activation to the threat level posed by the invader. For rapidly replicating invaders such as lytic phages that pose a major threat, target transcripts will accumulate in the cell, crossing the threshold and induce a robust immune response. For invaders such as lysogenic phages or mobile plasmids that take up residence in the cell and thus pose a minor threat, target transcripts may be expressed at a sufficiently low level to not induce the immune response. If targeting leads to gene silencing of essential transcripts, it could block invader replication, subsequently clearing the invader while sparing the infected cell from dormancy. In this setup, induced dormancy would instead represent a back-up strategy if the target transcript continues to accumulate in the cell.

For invader defense, one ramification of the target expression threshold is that an Acr (Bondy-Denomy et al., 2013; Marino et al., 2020) that reduces the concentration of activated Cas13a complexes could rapidly interfere with immune activation. Accordingly, recent work reporting one of the first Acrs against Cas13a showed that the invading Acr-encoding phage could shut down CRISPR defenses and proliferate in the first wave of infection (Meeske et al., 2020). In contrast, Acrs that block DNA-targeting systems could allow only a second wave of infecting phages to proliferate, as the defenses would clear the first wave before the Acrs could fully inhibit the CRISPR defenses (Borges et al., 2018; Landsberger et al., 2018). The prior work on the Cas13a Acr attributed escape by the Acr-encoding phage to Cas13a degrading the phage transcripts but not the phage DNA, allowing the phage to eventually recover. However, we found that lowering the concentration of activated Cas13a complexes could allow phage transcripts to cross the original expression threshold without inducing dormancy. Therefore, both mechanisms could be at work for phages encoding Cas13 Acrs.

Beyond Cas13a, Type III CRISPR-Cas systems also target RNA and can elicit collateral RNA cleavage that could also be impacted by a target expression threshold (Kazlauskiene et al., 2017; Niewoehner et al., 2017; Rostøl and Marraffini, 2019). For these systems, collateral RNA cleavage is induced through Cas10 within the activated effector complex synthesizing cyclic oligo adenylates (cOAs) from ATP. These small molecules then bind to different accessory proteins such as Csm6/Csx1 encoded by some Type III CRISPR-Cas systems, which then begin non-specifically cleaving cellular RNAs. A key difference compared to Cas13 is that one activated Type III effector complex can produce a large number of cOA molecules, potentially activating a large number of accessory proteins. This intermediate amplification step could effectively eliminate the threshold, leading to widespread collateral RNA cleavage and dormancy. However, depending on how quickly the cOAs are synthesized, diffuse, and turnover, a minimal threshold of activated effector complexes may be necessary to elicit widespread RNA cleavage. Furthermore, the existence of cOA-degrading anti-CRISPR proteins could offer a means to artificially raise the target expression threshold to the point where this mode of immunity is never activated over the course of the infection. Similar behaviors may be expected for Type VI CRISPR-Cas systems encoding Cas13b and the accessory protein Csx28 that appears to amplify the immune response (Smargon et al., 2017; VanderWal et al., 2021). Exploring the extent to which a target expression threshold exists for different Type III and VI-B CRISPR-Cas systems could broaden our understanding of the target expression threshold, particularly when back-up defenses exist that enhance the immune response.

Finally, the target expression threshold may help resolve otherwise contradictory observations when implementing Cas13-based technologies in bacteria and in eukaryotic cells. In bacteria, Cas13 was previously thought to induce widespread RNA cleavage as long as the target transcript was present (Abudayyeh et al., 2016). In contrast, in eukaryotic cells, Cas13 was reported to function as a sequence-specific gene silencer with no obvious off-target effects (Abudayyeh et al., 2017; Huynh et al.; Konermann et al., 2018; Mahas et al., 2019). To resolve this clear discrepancy, Cas13 was proposed to function differently in bacteria and in eukaryotic cells. At the same time, there are emerging reports of Cas13 inducing collateral RNA cleavage in human cells, although cell-type specific factors were suggested as the underlying cause (Özcan et al., 2021; Wang et al., 2019; Wu et al., 2021). The target expression threshold helps unify these observations. In bacteria, target transcripts under threshold do not induce widespread collateral RNA cleavage and instead are unaffected by targeting or undergo silencing. Applying our proposed biochemical mechanism to eukaryotic cells, activated Cas13 complexes would be much less concentrated given the larger size of the cytoplasm and the distribution of transcripts in this cellular compartment that lend to a higher expression threshold. The few examples of collateral cleavage might represent scenarios where the concentration of activated Cas13 complexes or the target transcript is much higher or localized close to important cellular RNAs. Investigating Cas13 nucleases that exhibit less collateral activity (Abudayyeh et al., 2017; Xu et al., 2021) could pose a simple solution to avoid potentially catastrophic RNA cleavage. The target expression threshold therefore may impact not only bacterial immune defense but also the application of Cas13 for programmable gene silencing in eukaryotic cells as well as for antivirals (Abbott et al., 2020; Abudayyeh et al., 2017; Konermann et al., 2018).

## STAR METHODS

### Strains and plasmids

The strains and plasmids used in this study are listed in Table S2. The main plasmids used are pZ003 (LshCas13a), pFT50 (crRNA backbone), pFT50-repeat (crRNA backbone with repeat) and pFT62 (target backbone). Spacers were inserted into the backbone by digestion with BsmBI and ligation with Instant sticky-end ligase master mix (NEB, M0370S). Q5 mutagenesis or Gibson assembly were used to modify the target plasmid or the nuclease backbone. All of the oligos used in this research can be found in Table S3.

### Transformation-based targeting assays

The transformation assays were conducted in two ways: targeting a genomically-encoded transcript and a plasmid-encoded transcript. For genomically-encoded transcripts, biological replicates containing the nuclease plasmid were inoculated overnight in LB medium (10 g tryptone, 5 g yeast extract, and 10 g NaCl in 1 L of dH_2_O) with chloramphenicol (Cm, 34 μg/mL). After 16 h, the ABS_600_ was measured and the samples were normalized, back-diluted 1:50 in fresh LB with Cm, and grown until an ABS_600_ of 0.6 - 0.8. Cultures were placed on ice and made electrocompetent by washing the pellet twice with 10% glycerol. Then, 50 ng of the crRNA plasmids were transformed into 40 μL of competent cells using the *E. coli* 1 program on the MicroPulser Electroporator (Bio-rad). After 1 h of recovery in 500 μL of SOC medium (SOB medium: 20 g Tryptone, 5 g Yeast Extract, 0.5 g NaCl, 800 mL dH_2_O and 10 mL 250 mM KCl adjusted to pH 7. To SOB medium added 5ml 2M MgCl_2_, 20 ml of 1 M Glucose), 10-fold dilutions of the cultures in 1X PBS (10X PBS: 80 g NaCL, 2 g KCl, 17,7 g Na_2_HPO_4_*2 H_2_O, 2.72 g KH_2_PO_4_, fill up to 1 L with mqH_2_O, set pH to 7.4 and autoclave) were prepared and 5 μL spot dilutions were plated on Cm and ampicillin (Amp, 100 μg/mL) LB plates and incubated at 37°C for 16-18 h.

For the plasmid-encoded transcripts, cultures containing the nuclease plasmid and the target plasmid were transformed with the crRNA plasmid. The rest of the procedure matched that followed for the genome targeting assay. The synthetic sequence expressed on a plasmid (pFT62) contains part of the mRFP1 gene sequence, which does not match the *E. coli* genome. Fragments from mRNAs (target plus 20 bp upstream and downstream) were cloned in pFT62 in place of the synthetic target. Finally, to see if cloning an entire gene on a plasmid would change the targeting outcome, full gene sequences with RBS and stop codon have been cloned in pFT62 through Gibson assembly.

### Flow cytometry analysis

Plasmids expressing GFP under different Anderson promoters were cloned by Q5 mutagenesis to introduce the different promoters in the pUA66-PJ23119 GFP plasmid. Cells containing the nuclease plasmid and the GFP plasmid were inoculated overnight in LB medium with kanamycin (Kan, 50 μg/mL) and Cm, then normalized, back-diluted to ABS_600_ = 0.02 and cultured until ABS_600_ ≈ 0.8. Cells were then pelleted and resuspended in 1X PBS before being applied to an Accuri C6 Plus analytical flow cytometer (BD Biosciences). 30,000 events were obtained by gating on living cells, and the mean of the FL1-H values were quantified as GFP fluorescence. Final fluorescence values were obtained by subtracting the autofluorescence of cells not expressing GFP.

### L-arabinose induction of Cas13a-mediated targeting in *E. coli*

The plasmid with the synthetic target under control of the arabinose-inducible promoter was cloned by introducing the P_BAD_ promoter in pFT62 through Gibson assembly. To validate this system, *E. coli* MG1655 Δ*araBAD* P_con_-*araFGH* cells were transformed with the nuclease, the gRNA and the arabinose-inducible target plasmids. Three biological replicates were inoculated overnight in LB supplemented with Amp, Cm, and Kan as well as 0.2% glucose to reduce background expression from the P_BAD_ promoter. The samples were then pelleted, resuspended in LB with antibiotics and then back-diluted to ABS_600_ = 0.01 with or without 0.2% L-arabinose. Culture turbidity was recorded over time on a Synergy Neo2 or H1 fluorescence microplate reader (BioTek) for 16 h by measuring ABS_600_ every 3 min.

### Assessment of collateral RNA cleavage

Using the validated arabinose-inducible setup, collateral RNA cleavage was visualized on an agarose gel. Cells with the nuclease, gRNA, and target plasmids were grown overnight in Cm, Amp, Kan LB with 0.2% glucose, washed in LB with antibiotics to remove glucose, and back-diluted to ABS_600_ = 0.01. The cells were cultured until ABS_600_ ≈ 0.4, after which each sample was split in equal volumes with or without the inducer. After 1 h from induction, the ABS_600_ of each sample was measured, and the same number of cells (2250 ABS_600_×mL) was snap frozen on dry ice. The next day, total RNA was extracted using the Directzol RNA mini-prep plus kit (Zymo research). Then 1 μg of each RNA was mixed 2:3 with an RNA loading dye (2x) (for 50 mL: 625 µL Bromophenol blue 2%, 625 µL 2% Xylene Cyanol, 1800 µL of 0.5 M EDTA pH = 8.0, 46,821 mL Formamide), heated at 70°C for 10 min, placed on ice, and resolved on a 1% TBE gel at 120 V for 40 min. RiboRuler High Range RNA ladder (Thermo Scientific, SM1821) was used as a size marker.

### Library design and validation

The reference genome and annotation of *E. coli* K12 MG1655 (NC_000913.3) was used for gRNA library design. First, all potential 32-nt guides were designed for protein-coding genes (limited to the CDS) and rRNAs with non-GU PFS and GC content between 40% and 60%, resulting in an average of 484 guides per gene. To reduce the size of the library, and considering the unknown effect of the targeting location within a gene on guide efficiency, each gene was divided into a maximum of 10 sections with equal length. Within each section, guides were filtered based on the strength of local secondary structure, defined as ΔG, in both repeat-guide sequence and the mRNA targeting region (including a region of 2 times length of gRNA before and after the target). ΔG was calculated as the energy difference between the unconstrained minimum free energy (MFE) structure and the constrained MFE structure with no base pairs, estimated using RNAfold from the Vienna RNA Package (Lorenz et al., 2011) version 2.4.12. Sequences (either guide or flanking primer sequences) containing BsmBI restriction sites or homopolymer stretches of more than four consecutive nucleotides were excluded to facilitate synthesis and cloning. The guide with the lowest secondary structure strength in each section was selected, resulting in a library of 25,997 guides, including 25,470 guides targeting protein-coding genes, 127 guides targeting rRNAs, and 400 randomized non-targeting guides as negative controls. A sequence containing a universal primer binding site and the BsmBI restriction site was added to the guides to amplify the oligo library and digest it before ligating it into the backbone (see Table S4). The library was synthesized by Twist Bioscience.

The base backbone pFT50 was slightly modified to insert the direct repeat before the GFP dropout site to be able to limit the insertion size to the spacer itself. A BsmBI restriction site present in pFT50 was also eliminated through Q5 mutagenesis.

### Guide library cloning and verification

The library was amplified with Kapa Hifi polymerase (20 ng DNA) for 10 cycles following the manufacturer’s instructions (Ta = 64°C; 30 s denaturation, 20 s annealing, 15 s extension) using primers SPCpr 349/350. 5 µL library (150 nM) and 5 µL of backbone (50 nM) were mixed in 25 µL of total reaction volume. The mixture was subjected to 50 cycles of BsmBI digestion (3 min at 42°C) and ligation (T4 ligase - 5 min at 16°C) with a final digestion at 55°C for 60 min to ensure complete removal of the backbone, followed by a 10 minute heat inactivation at 80°C. The sample was then ethanol precipitated, and 5 µg were transformed into fresh electrocompetent Top10 cells (90 µL). The transformation was conducted with two separate batches of electrocompetent cells to ensure enough transformants were obtained. After recovering the two cultures in 500µL of SOC medium shaking at 37°C for 1 h, the recovered cultures were back-diluted into 150 mL LB with Amp and cultured with shaking at 37°C for 12 h. The next day, plasmid DNA from the culture was isolated using the ZymoPURE II Maxiprep Kit (Zymo Research, D4203) and further purified by ethanol precipitation.

### Guide library screen

Two replicates of *E. coli* MG1655 cells with or without the nuclease plasmid were inoculated overnight, then the next day the ABS_600_ was normalized and the cells back-diluted to ABS_600_ ≈ 0.1 in fresh LB with or without Cm. Once each culture reached ABS_600_ ≈ 0.8, cells were made electrocompetent by washing twice with 10% glycerol and finally resuspended in 480 μL 10% glycerol. For each sample, six separate transformations were conducted each with 1 μg of library DNA (40 μL/transformation). Transformed cells were then recovered in 500 μL SOC medium for 1 h with shaking at 37°C. The six reactions were combined to yield 3 mL of culture per condition. Serial dilutions of this culture were made, and 100 μL of 1:10,000 dilutions were plated for the targeting and no-Cas13a samples with the appropriate antibiotics (Cm and Amp, or Amp only), yielding a theoretical library coverage of ∼9,500. The remaining culture was diluted 1:100 in LB with Amp and Cm to ABS_600_ = 0.06 and cultured for 12 h with shaking at 37°C. Finally, the library was isolated with the ZymoPure II Plasmid Midiprep Kit (Zymo Research, D4200).

### Next-generation sequencing of the guide libraries

The guides sequences from the purified library DNA were amplified with Kapa Hifi polymerase using primers oEV-315/316 (NT1), oEV-317/318 (NT2), oEV-319/320 (T1), oEV-321/322 (T2). 10 ng of DNA were included in a 50-μL PCR reaction for 15 amplification cycles (15 s at 98°C, 30 s at 64°C, 30 s extension). The amplification products were purified using Ampure beads and further amplified with primers oEV-323/324 (NT1), oEV-325/326 (NT2), oEV-327/328 (T1), oEV-329/330 (T2) to add the appropriate indices and Illumina adaptors. For this reaction, the same settings were used with the only difference being the amount of input DNA (25 ng) and the number of cycles (10). The resulting amplification products were purified with Ampure beads and resolved on a gel to verify the presence of the correct amplicon. The samples were submitted for Sanger sequencing and Bioanalyzer analysis as a quality check. Finally, samples were submitted for next-generation sequencing at the NextSeq 500 sequencer (Illumina) with a 150 bp paired-ends kit (130 million reads) to obtain 1000-fold coverage. To increase the library diversity, 20% of phiX phage was spiked-in.

To correlate transcript expression levels with guide depletion, we measured transcript levels in *E. coli* MG1655 under conditions paralleling the library screen. Briefly, two replicates of cells containing the plasmid cBAD33 (empty backbone for nuclease plasmid) were cultured overnight and then normalized to ABS_600_ = 0.06 in LB with Cm and cultured to ABS_600_ ≈ 0.5 or ABS_600_ ≈ 0.8. At those growth points, cells were pelleted and snap-frozen for RNA extraction with the Directzol RNA mini-prep plus kit (Zymo Research, R2071). The samples were also DNase-treated with TURBO DNase (Thermo Fisher Scientific, AM2239) and quality verified using a Bioanalyzer 2100 (Agilent). Finally, rRNA was removed with the Rybo-off rRNA depletion kit (Vazyme Biotech, N407-01) and the samples were sequenced on a NovaSeq 6000 (Illumina) with 50-bp paired-end reads.

The resulting NGS data were deposited in NCBI’s Gene Expression Omnibus (Edgar et al., 2002) and are accessible through GEO Series accession number GSE179913 for the genome-wide screen (https://www.ncbi.nlm.nih.gov/geo/query/acc.cgi?acc=GSE179913) and GSE179914 for transcriptomic analysis (https://www.ncbi.nlm.nih.gov/geo/query/acc.cgi?acc=GSE179914).

### Screen analysis and machine learning model

For analysis of the genome-wide screen, after merging using BBMerge (version 38.69) with parameters “qtrim2=t, ecco, trimq=20, -Xmx1g, mix=f”, paired-end sequence reads with a perfect match were assigned to gRNA sequences. After filtering guides for at least 1 count per million reads in at least 2 samples, the library sizes were normalized using the read counts for non-targeting guides with the trimmed mean of M-values (TMM) method in edgeR (Robinson and Oshlack, 2010; Robinson et al., 2009) (version 3.28.0). Differential abundance of gRNAs between targeting samples and control samples lacking the Cas13a nuclease was assessed using the edgeR quasi-likelihood F test after fitting a generalized linear model. The translation initiation rate of each gene was predicted using RBS calculator (version 1.0) (Salis, 2011).

For RNA seq analysis, sequencing reads were aligned to the *E. coli* K12 MG1655 genome (NC_000913.3) using STAR (Dobin et al., 2013) (version 2.7.4a) with parameters “--alignIntronMax 1 --genomeSAindexNbases 10 --outSAMtype BAM SortedByCoordinate” and the count of reads mapping to each gene was obtained using HTSeq (Anders et al., 2015) (version 0.9.1) with parameters “-i locus_tag -r pos --stranded reverse --nonunique none -t gene”, followed by calculating transcripts per million (TPM).

The machine learning regression model was developed with 144 features as predictors and the log2FC values of gRNAs from the genome-wide screen as targets using auto-sklearn version 0.10.0 (Feurer et al., 2019) with all possible estimators and preprocessors included and parameters ’ensemble_size’: 1, ’resampling_strategy’: ’cv’, ’resampling_strategy_arguments’: {’folds’: 5}, ’per_run_time_limit’: 360, and ’time_left_for_this_task’: 3600. Features included gene expression level (log2 transformed TPM at OD 0.4), gene essentiality, gene id, gene length, (percent) targeting position in gene, delta G of repeat-gRNA and mRNA targeting region, and the one-hot-encoded PFS sequence and gRNA sequence. Gene essentiality information in LB Lennox medium was obtained from EcoCyc (Keseler et al., 2017) (https://ecocyc.org). The optimal histogram-based gradient boosting model was evaluated using 10-fold cross-validation and interpreted using TreeSHAP version 0.36.0 (Lundberg et al., 2020).

### Targeting assay with lower nuclease expression

In order to look at targeting with varied nuclease expression, we used a plasmid expressing the nuclease under the constitutive promoter PJ23108 (Pw). At first we assessed the relative promoter strength of Pw and the native promoter (P_native_) by cloning the *gfp* gene in the nuclease backbone in place of the nuclease itself through Gibson assembly. Each *gfp* plasmid was transformed with the target plasmid into *E. coli* MG1655 cells, and the ‘Flow cytometry analysis’ protocol was followed to measure fluorescence of the two constructs. We then performed the transformation-based targeting assay with the low expressed nuclease, the crRNA (T1) and different Anderson promoters (P1 to P8) cloned in front of the synthetic target sequence.

### Cell-free transcription-translation assays

Plasmids encoding the nuclease, a crRNA, a target sequence, and deGFP were used to assess the targeting activity in a cell-free transcription-translation assay. The nuclease was either under the control of P_native_ or Pw and the crRNA encoded either a targeting (T) spacer or non-targeting (NT) control. To assess differences in targeting activity when the nuclease is under the control of different promoters, the two nuclease constructs were added separately with either the targeting or the non-targeting crRNA and the targeted plasmid to *myTXTL* Sigma 70 Master Mix (Arbor Biosciences, 507005), with a final concentration of 2 nM, 1 nM and 0.5 nM, respectively. The samples were incubated for 2 h at 29°C, before a plasmid encoding deGFP was added to a final concentration of 0.5 nM. The samples were then incubated at 29°C for 16 h in a plate reader (BioTek Synergy Neo2) and fluorescence was measured every three minutes (excitation, emission: 485 nm, 528 nm). All shown data was produced using the Echo 525 Liquid Handler (Beckman Coulter). The assays were therefore scaled down to 3 µl reactions per replicate, with four replicates each. As part of the analysis, the background fluorescence from *myTXTL* mix and water samples was subtracted from all samples. Grubb’s test was performed using the values after 16 h to identify outliers between replicates (α = 0.1). If no outliers were identified, the first of the four replicates was discarded. The graph shows the average deGFP fluorescence over time together with the standard deviation.

### Infection experiments with MS2 phage

MS2 phage concentration (PFU/mL) has been calculated by performing a plaque assay. Transcription-based targeting assays have been performed by transforming *E. coli* CGSC 4401 cells (F+) expressing the nuclease and differentially abundant target *rep* or *cp* gene transcripts with the correspondent targeting guides and by calculating the reduction in colony numbers compared to a non-targeting guide. *E. coli* cells containing the active or dead nuclease and the guide encoding plasmids have been grown overnight in selective LB medium, back-diluted to ABS_600_ = 0.05 and let grow until ABS_600_ ≈ 0.3. The samples have then been normalized to ABS_600_ = 0.3 and aliquoted in a 96 well plate (Thermo Scientific, 167008) together with different amounts of phages (MOI 0.1 and 5) or the respective volume of LB for the no-infection control. Cell growth has been recorded over 16 h by measuring ABS_600_ every three minutes in a plate reader at 37°C.

### Tolerance to targeted *kanR* gene

*E. coli* colonies containing the nuclease and crRNA (K1, K2) plasmids were transformed with the targeted plasmid, which has a cloDF13 ori, different promoters in front of the *kanR* gene (P2, P5, P8), and a non-targeted hygromycin (Hyg, 100 μg/mL) resistance cassette. The plasmid was cloned in two steps by Gibson assembly with pUA66 as the backbone. K1 and K2 were targeting two different regions within the *kanR* transcript. The procedure is analogous to the transformation assay with a plasmid-encoded transcript, with the targeted plasmid selected on Hyg. The next day, colonies from the spot dilutions were counted, and colonies from the samples showing a negligible reduction in transformation compared to the non-targeting control were inoculated overnight in LB with Cm, Amp and Hyg. Then the cultures were washed to remove Hyg and back-diluted to ABS_600_ = 0.01 in either Hyg or Kan. The growth curves for the different conditions were measured over time on the microplate reader for 14 h at 37°C by measuring ABS_600_ every 3 min. For generating the heatmap, 12 h time points were selected to compare the different conditions and the resulting graph is the average of 9 biological replicates.

### Data analysis and image visualization

Microsoft Excel was used to analyze the data, and GraphPad Prism was used to generate the bar plots and heatmaps. The graphs were then modified in Adobe Illustrator to construct the final figures. Transformation fold-reduction in the transformation assays was calculated as the ratio between non-targeting and targeting colonies.

### Statistical analyses

All statistical analyses were performed using a Welch’s t test assuming unequal variances. P-values above 0.05 or average values lower than the reference average were considered non-significant. Statistical comparisons for the transformation assays relied on log values, which assumes the samples are normally distributed on a log scale.

## ACKNOWLEDGMENTS

We thank Fani Ttofali and Chunyu Liao for providing initial plasmids from which constructs used in this work were generated. We also thank Tatjana Achmedov for technical advice and assistance. This work was supported by the Joint Programming Initiative on Antimicrobial Resistance (01KI1824 to C.L.B.), the National Institutes of Health (1R35GM119561 to C.L.B.), a bayresq.net Bavarian research network grant (to L.B.), and the Defense Advanced Research Projects Agency Safe Genes programme (HR0011-17-2-0042 to C.L.B.). The views, opinions and/or findings expressed should not be interpreted as representing the official views or policies of the Department of Defense or the U.S. Government.

## AUTHOR CONTRIBUTIONS

Conceptualization: E.V., C.L.B.; Methodology: E.V., Y.Y., L.B., C.L.B.; Software: Y.Y., L.B.; Validation: E.V., Y.Y., S.P.C., K.G.W.; Formal analysis: E.V., Y.Y., K.G.W., L.B.; Investigation: E.V., S.P.C., K.G.W.; Writing - Original Draft: E.V., Y.Y., L.B., C.L.B.; Writing - Review and Editing: E.V., Y.Y., S.P.C., K.G.W., L.B., C.L.B; Visualization: E.V., Y.Y., K.G.W., C.L.B.; Supervision: L.B., C.L.B; Funding acquisition: L.B., C.L.B.

## REFERENCES

1. Abbott, T.R., Dhamdhere, G., Liu, Y., Lin, X., Goudy, L., Zeng, L., Chemparathy, A., Chmura, S., Heaton, N.S., Debs, R., et al. (2020). Development of CRISPR as an Antiviral Strategy to Combat SARS-CoV-2 and Influenza. Cell 181, 865–876.e12.

2. Abudayyeh, O.O., Gootenberg, J.S., Konermann, S., Joung, J., Slaymaker, I.M., Cox, D.B.T., Shmakov, S., Makarova, K.S., Semenova, E., Minakhin, L., et al. (2016). C2c2 is a single-component programmable RNA-guided RNA-targeting CRISPR effector. Science 353, aaf5573.

3. Abudayyeh, O.O., Gootenberg, J.S., Essletzbichler, P., Han, S., Joung, J., Belanto, J.J., Verdine, V., Cox, D.B.T., Kellner, M.J., Regev, A., et al. (2017). RNA targeting with CRISPR-Cas13. Nature 550, 280–284.

4. Anders, S., Pyl, P.T., and Huber, W. (2015). HTSeq--a Python framework to work with high-throughput sequencing data. Bioinformatics 31, 166–169.

5. Barrangou, R., Fremaux, C., Deveau, H., Richards, M., Boyaval, P., Moineau, S., Romero, D.A., and Horvath, P. (2007). CRISPR provides acquired resistance against viruses in prokaryotes. Science 315, 1709–1712.

6. Bondy-Denomy, J., Pawluk, A., Maxwell, K.L., and Davidson, A.R. (2013). Bacteriophage genes that inactivate the CRISPR/Cas bacterial immune system. Nature 493, 429–432.

7. Borges, A.L., Zhang, J.Y., Rollins, M.F., Osuna, B.A., Wiedenheft, B., and Bondy-Denomy, J. (2018). Bacteriophage Cooperation Suppresses CRISPR-Cas3 and Cas9 Immunity. Cell 174, 917–925.e10.

8. Buchman, A.B., Brogan, D.J., Sun, R., Yang, T., Hsu, P.D., and Akbari, O.S. (2020). Programmable RNA Targeting Using CasRx in Flies. CRISPR J 3, 164–176.

9. Charpentier, E., Richter, H., van der Oost, J., and White, M.F. (2015). Biogenesis pathways of RNA guides in archaeal and bacterial CRISPR-Cas adaptive immunity. FEMS Microbiol. Rev. 39, 428–441.

10. Cox, D.B.T., Gootenberg, J.S., Abudayyeh, O.O., Franklin, B., Kellner, M.J., Joung, J., and Zhang, F. (2017). RNA editing with CRISPR-Cas13. Science 358, 1019–1027.

11. Dobin, A., Davis, C.A., Schlesinger, F., Drenkow, J., Zaleski, C., Jha, S., Batut, P., Chaisson, M., and Gingeras, T.R. (2013). STAR: ultrafast universal RNA-seq aligner. Bioinformatics 29, 15–21.

12. East-Seletsky, A., O’Connell, M.R., Knight, S.C., Burstein, D., Cate, J.H.D., Tjian, R., and Doudna, J.A. (2016). Two distinct RNase activities of CRISPR-C2c2 enable guide-RNA processing and RNA detection. Nature 538, 270–273.

13. Elowitz, M.B., Levine, A.J., Siggia, E.D., and Swain, P.S. (2002). Stochastic Gene Expression in a Single Cell. Science 297, 1183–1186.

14. Feurer, M., Klein, A., Eggensperger, K., Springenberg, J.T., Blum, M., and Hutter, F. (2019). Auto-sklearn: Efficient and Robust Automated Machine Learning. Automated Machine Learning 113–134.

15. Frost, L.S., Leplae, R., Summers, A.O., and Toussaint, A. (2005). Mobile genetic elements: the agents of open source evolution. Nat. Rev. Microbiol. 3, 722–732.

16. Gasiunas, G., Barrangou, R., Horvath, P., and Siksnys, V. (2012). Cas9-crRNA ribonucleoprotein complex mediates specific DNA cleavage for adaptive immunity in bacteria. Proc. Natl. Acad. Sci. U. S. A. 109, E2579–E2586.

17. Goldberg, G.W., and Marraffini, L.A. (2015). Resistance and tolerance to foreign elements by prokaryotic immune systems — curating the genome. Nature Reviews Immunology 15, 717– 724.

18. Hampton, H.G., Watson, B.N.J., and Fineran, P.C. (2020). The arms race between bacteria and their phage foes. Nature 577, 327–336.

19. Hille, F., Richter, H., Wong, S.P., Bratovič, M., Ressel, S., and Charpentier, E. (2018). The Biology of CRISPR-Cas: Backward and Forward. Cell 172, 1239–1259.

20. Huynh, N., Depner, N., Larson, R., and King-Jones, K. A versatile toolkit for CRISPR-Cas13-based RNA manipulation in Drosophila.

21. Jackson, S.A., McKenzie, R.E., Fagerlund, R.D., Kieper, S.N., Fineran, P.C., and Brouns, S.J.J. (2017). CRISPR-Cas: Adapting to change. Science 356.

22. Jinek, M., Chylinski, K., Fonfara, I., Hauer, M., Doudna, J.A., and Charpentier, E. (2012). A programmable dual-RNA-guided DNA endonuclease in adaptive bacterial immunity. Science 337, 816–821.

23. Kazlauskiene, M., Kostiuk, G., Venclovas, Č., Tamulaitis, G., and Siksnys, V. (2017). A cyclic oligonucleotide signaling pathway in type III CRISPR-Cas systems. Science 357, 605–609.

24. Keseler, I.M., Mackie, A., Santos-Zavaleta, A., Billington, R., Bonavides-Martínez, C., Caspi, R., Fulcher, C., Gama-Castro, S., Kothari, A., Krummenacker, M., et al. (2017). The EcoCyc database: reflecting new knowledge about Escherichia coli K-12. Nucleic Acids Res. 45, D543– D550.

25. Kiga, K., Tan, X.-E., Ibarra-Chávez, R., Watanabe, S., Aiba, Y., Sato’o, Y., Li, F.-Y., Sasahara, T., Cui, B., Kawauchi, M., et al. (2020). Development of CRISPR-Cas13a-based antimicrobials capable of sequence-specific killing of target bacteria. Nat. Commun. 11, 2934.

26. Konermann, S., Lotfy, P., Brideau, N.J., Oki, J., Shokhirev, M.N., and Hsu, P.D. (2018). Transcriptome Engineering with RNA-Targeting Type VI-D CRISPR Effectors. Cell 173, 665– 676.e14.

27. Koonin, E.V., Makarova, K.S., Wolf, Y.I., and Krupovic, M. (2020). Evolutionary entanglement of mobile genetic elements and host defence systems: guns for hire. Nat. Rev. Genet. 21, 119– 131.

28. Landsberger, M., Gandon, S., Meaden, S., Rollie, C., Chevallereau, A., Chabas, H., Buckling, A., Westra, E.R., and van Houte, S. (2018). Anti-CRISPR Phages Cooperate to Overcome CRISPR-Cas Immunity. Cell 174, 908–916.e12.

29. Leenay, R.T., and Beisel, C.L. (2017). Deciphering, Communicating, and Engineering the CRISPR PAM. J. Mol. Biol. 429, 177–191.

30. Liao, C., Ttofali, F., Slotkowski, R.A., Denny, S.R., Cecil, T.D., Leenay, R.T., Keung, A.J., and Beisel, C.L. (2019). Modular one-pot assembly of CRISPR arrays enables library generation and reveals factors influencing crRNA biogenesis. Nat. Commun. 10, 2948.

31. Liu, L., Li, X., Wang, J., Wang, M., Chen, P., Yin, M., Li, J., Sheng, G., and Wang, Y. (2017a). Two Distant Catalytic Sites Are Responsible for C2c2 RNase Activities. Cell 168, 121–134.e12.

32. Liu, L., Li, X., Ma, J., Li, Z., You, L., Wang, J., Wang, M., Zhang, X., and Wang, Y. (2017b). The Molecular Architecture for RNA-Guided RNA Cleavage by Cas13a. Cell 170, 714–726.e10.

33. Lundberg, S.M., Erion, G., Chen, H., DeGrave, A., Prutkin, J.M., Nair, B., Katz, R., Himmelfarb, J., Bansal, N., and Lee, S.-I. (2020). From local explanations to global understanding with explainable AI for trees. Nature Machine Intelligence 2, 56–67.

34. Mahas, A., Aman, R., and Mahfouz, M. (2019). CRISPR-Cas13d mediates robust RNA virus interference in plants. Genome Biol. 20, 263.

35. Makarova, K.S., Wolf, Y.I., Alkhnbashi, O.S., Costa, F., Shah, S.A., Saunders, S.J., Barrangou, R., Brouns, S.J.J., Charpentier, E., Haft, D.H., et al. (2015). An updated evolutionary classification of CRISPR-Cas systems. Nat. Rev. Microbiol. 13, 722–736.

36. Makarova, K.S., Wolf, Y.I., Iranzo, J., Shmakov, S.A., Alkhnbashi, O.S., Brouns, S.J.J., Charpentier, E., Cheng, D., Haft, D.H., Horvath, P., et al. (2020). Evolutionary classification of CRISPR-Cas systems: a burst of class 2 and derived variants. Nat. Rev. Microbiol. 18, 67–83.

37. Marino, N.D., Pinilla-Redondo, R., Csörgő, B., and Bondy-Denomy, J. (2020). Anti-CRISPR protein applications: natural brakes for CRISPR-Cas technologies. Nature Methods 17, 471– 479.

38. Marraffini, L.A., and Sontheimer, E.J. (2010). CRISPR interference: RNA-directed adaptive immunity in bacteria and archaea. Nat. Rev. Genet. 11, 181–190.

39. Marshall, R., Maxwell, C.S., Collins, S.P., Jacobsen, T., Luo, M.L., Begemann, M.B., Gray, B.N., January, E., Singer, A., He, Y., et al. (2018). Rapid and Scalable Characterization of CRISPR Technologies Using an E. coli Cell-Free Transcription-Translation System. Mol. Cell 69, 146– 157.e3.

40. Marshall, R., Beisel, C.L., and Noireaux, V. (2020). Rapid Testing of CRISPR Nucleases and Guide RNAs in an Cell-Free Transcription-Translation System. STAR Protoc 1, 100003.

41. Meeske, A.J., and Marraffini, L.A. (2018). RNA Guide Complementarity Prevents Self-Targeting in Type VI CRISPR Systems. Mol. Cell 71, 791–801.e3.

42. Meeske, A.J., Nakandakari-Higa, S., and Marraffini, L.A. (2019). Cas13-induced cellular dormancy prevents the rise of CRISPR-resistant bacteriophage. Nature 570, 241–245.

43. Meeske, A.J., Jia, N., Cassel, A.K., Kozlova, A., Liao, J., Wiedmann, M., Patel, D.J., and Marraffini, L.A. (2020). A phage-encoded anti-CRISPR enables complete evasion of type VI-A CRISPR-Cas immunity. Science 369, 54–59.

44. Niewoehner, O., Garcia-Doval, C., Rostøl, J.T., Berk, C., Schwede, F., Bigler, L., Hall, J., Marraffini, L.A., and Jinek, M. (2017). Type III CRISPR–Cas systems produce cyclic oligoadenylate second messengers. Nature 548, 543–548.

45. van der Oost, J., Westra, E.R., Jackson, R.N., and Wiedenheft, B. (2014). Unravelling the structural and mechanistic basis of CRISPR-Cas systems. Nat. Rev. Microbiol. 12, 479–492.

46. Özcan, A., Krajeski, R., Ioannidi, E., Lee, B., Gardner, A., Makarova, K.S., Koonin, E.V., Abudayyeh, O.O., and Gootenberg, J.S. (2021). Programmable RNA targeting with the single-protein CRISPR effector Cas7-11. Nature.

47. Partridge, S.R., Kwong, S.M., Firth, N., and Jensen, S.O. (2018). Mobile Genetic Elements Associated with Antimicrobial Resistance. Clinical Microbiology Reviews 31.

48. Ratner, H.K., Escalera-Maurer, A., Le Rhun, A., Jaggavarapu, S., Wozniak, J.E., Crispell, E.K., Charpentier, E., and Weiss, D.S. (2019). Catalytically Active Cas9 Mediates Transcriptional Interference to Facilitate Bacterial Virulence. Mol. Cell 75, 498–510.e5.

49. Robinson, M.D., and Oshlack, A. (2010). A scaling normalization method for differential expression analysis of RNA-seq data. Genome Biol. 11, R25.

50. Robinson, M.D., McCarthy, D.J., and Smyth, G.K. (2009). edgeR: A Bioconductor package for differential expression analysis of digital gene expression data. Bioinformatics 26, 139–140.

51. Rollie, C., Chevallereau, A., Watson, B.N.J., Chyou, T.-Y., Fradet, O., McLeod, I., Fineran, P.C., Brown, C.M., Gandon, S., and Westra, E.R. (2020). Targeting of temperate phages drives loss of type I CRISPR-Cas systems. Nature 578, 149–153.

52. Rostøl, J.T., and Marraffini, L.A. (2019). Non-specific degradation of transcripts promotes plasmid clearance during type III-A CRISPR-Cas immunity. Nat Microbiol 4, 656–662.

53. Salis, H.M. (2011). The ribosome binding site calculator. Methods Enzymol. 498, 19–42.

54. Smargon, A.A., Cox, D.B.T., Pyzocha, N.K., Zheng, K., Slaymaker, I.M., Gootenberg, J.S., Abudayyeh, O.A., Essletzbichler, P., Shmakov, S., Makarova, K.S., et al. (2017). Cas13b Is a Type VI-B CRISPR-Associated RNA-Guided RNase Differentially Regulated by Accessory Proteins Csx27 and Csx28. Mol. Cell 65, 618–630.e7.

55. Stern, A., Keren, L., Wurtzel, O., Amitai, G., and Sorek, R. (2010). Self-targeting by CRISPR: gene regulation or autoimmunity? Trends in Genetics 26, 335–340.

56. Theofilopoulos, A.N., Kono, D.H., and Baccala, R. (2017). The multiple pathways to autoimmunity. Nature Immunology 18, 716–724.

57. VanderWal, A.R., Park, J.-U., Polevoda, B., Kellogg, E.H., and O’Connell, M.R. (2021). CRISPR-Csx28 forms a Cas13b-activated membrane pore required for robust CRISPR-Cas adaptive immunity.

58. Wang, B., Zhang, T., Yin, J., Yu, Y., Xu, W., Ding, J., Patel, D.J., and Yang, H. (2021). Structural basis for self-cleavage prevention by tag:anti-tag pairing complementarity in type VI Cas13 CRISPR systems. Mol. Cell 81, 1100–1115.e5.

59. Wang, Q., Liu, X., Zhou, J., Yang, C., Wang, G., Tan, Y., Wu, Y., Zhang, S., Yi, K., and Kang, C. (2019). The CRISPR-Cas13a Gene-Editing System Induces Collateral Cleavage of RNA in Glioma Cells. Advanced Science 6, 1901299.

60. Watanabe, S., Cui, B., Kiga, K., Aiba, Y., Tan, X.-E., Sato’o, Y., Kawauchi, M., Boonsiri, T., Thitiananpakorn, K., Taki, Y., et al. (2019). Composition and Diversity of CRISPR-Cas13a Systems in the Genus. Front. Microbiol. 10, 2838.

61. Wessels, H.-H., Méndez-Mancilla, A., Guo, X., Legut, M., Daniloski, Z., and Sanjana, N.E. (2020). Massively parallel Cas13 screens reveal principles for guide RNA design. Nat. Biotechnol. 38, 722–727.

62. Wu, Y., Jin, W., Wang, Q., Zhou, J., Wang, Y., Tan, Y., Cui, X., Tong, F., Yang, E., Wang, J., et al. (2021). Precise editing of FGFR3-TACC3 fusion genes with CRISPR-Cas13a provides a personalized therapeutic strategy for the treatment of human glioblastoma. Mol. Ther.

63. Xu, C., Zhou, Y., Xiao, Q., He, B., Geng, G., Wang, Z., Cao, B., Dong, X., Bai, W., Wang, Y., et al. (2021). Programmable RNA editing with compact CRISPR–Cas13 systems from uncultivated microbes. Nature Methods 18, 499–506.

