## Supplementary information for "A target expression threshold dictates invader defense and autoimmunity by CRISPR-Cas13"

### SUPPLEMENTARY FIGURES

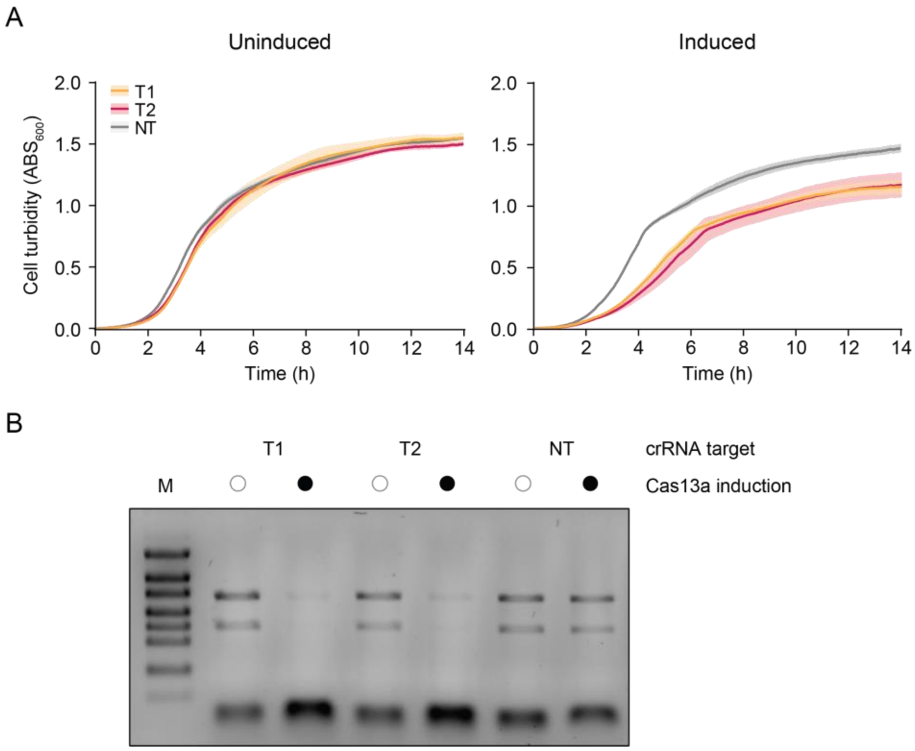


**Figure S1**. Cas13a targeting leads to inhibited growth and widespread RNA degradation, in line with its established role in inducing cellular dormancy. Related to **Fig. 1**.

(**A**) Inhibition of growth in liquid culture following induction of Cas13a-mediated targeting. The plasmid-encoded target is under the control of an arabinose-inducible promoter. *E. coli* cells are harboring the LshCas13a plasmid, the target plasmid, and the targeting (T1 or T2) or non-targeting (NT) crRNA plasmid. Back-diluted cells were supplemented with 0.2% arabinose (induced) or nothing (uninduced) at time 0 and cultured over time. Solid lines and light regions represent the mean and standard deviation of $ABS_{600}$ for three independent cultures.

(**B**) Impact of inducing target expression on total RNA. Total RNA was extracted from cultures after 1 h of induction and resolved on a 1% TBE gel. M: marker.

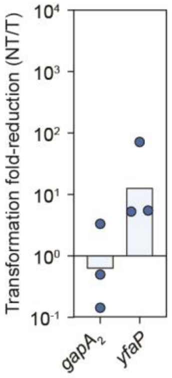


**Figure S2**. Impact of expressing the crRNA without Cas13a. Related to **Fig. 1**.

Genome targeting assay without nuclease reveals *yfaP* crRNA toxicity. The *gapA*$_2$ crRNA does not show a drop in colony numbers in the absence of LshCas13a. Fold-reduction was calculated as the ratio of transformants between the non-targeting (NT) and targeting (T) crRNA plasmids. The fold-reduction for the yfaP-targeting crRNA was similar to that observed with LshCas13a (see **Fig. 1C**), indicating that this crRNA is toxic when expressed in *E. coli* MG1655. Bars represent the geometric mean of triplicate independent experiments.

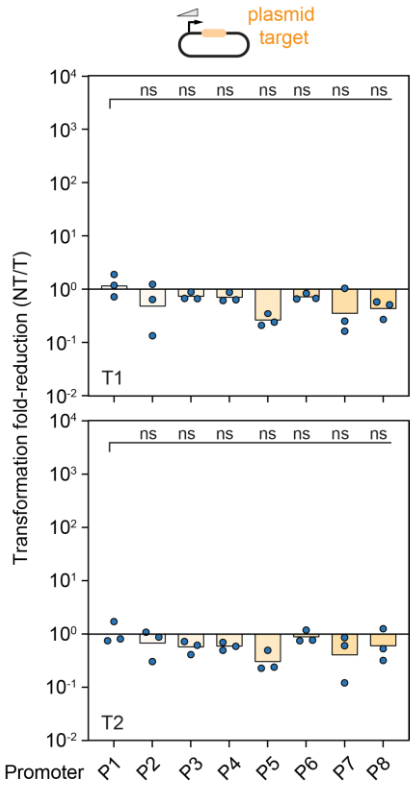


**Figure S3.** Impact of targeting the plasmid-encoded transcript expressed from different constitutive promoters by dCas13a. Related to **Fig. 2**.

Bars represent the geometric mean of triplicate independent experiments. Statistical significance was calculated by comparing the transformation fold-reduction to that of the weakest P1 promoter. ***: $p < 0.001$. **: $p < 0.01$. *: $p < 0.05$. ns: not significant or average below the reference.

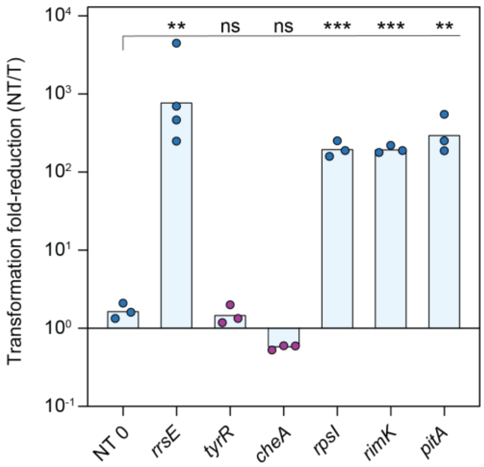


**Figure S4.** Selected highly-depleted guides from the library yield reduced transformation or growth. Related to **Fig. 3**.

A randomized crRNA from the library (NT 0) served as the negative control, while a crRNA targeting the rRNA encoded by *rrsE* served as a positive control. Purple data points represent small colonies indicative of reduced growth. Bars represent the mean of at least triplicate independent experiments. Statistical significance was calculated by comparing the transformation fold-reduction to that of NT 0. ***: $p < 0.001$. **: $p < 0.01$. *: $p < 0.05$. ns: not significant or average below the reference. Tested guides resulted in significant fold-reduction in transformation or small colonies.

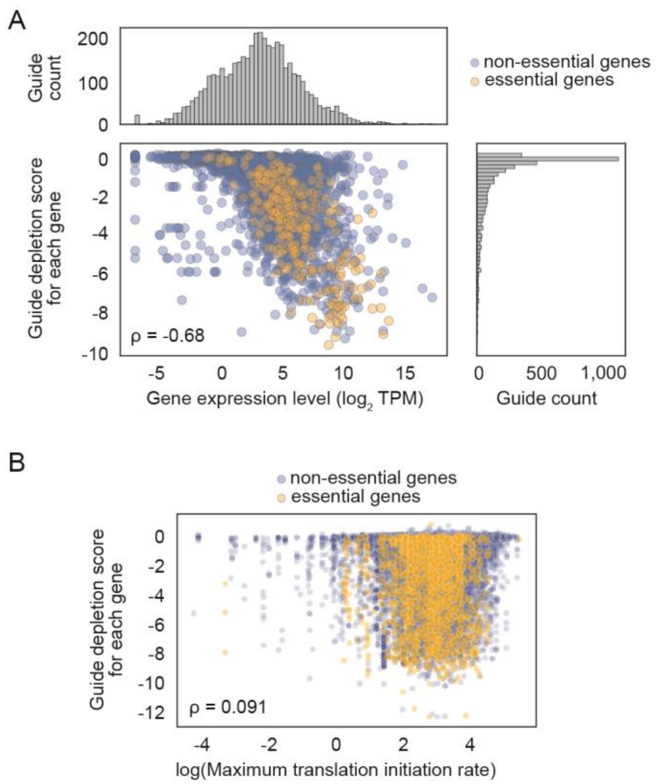


**Figure S5.** Additional results from the guide depletion screen. Related to **Fig. 3**.

(**A**) Correlation between guide depletion score and the expression levels of the target gene for cells cultured to a turbidity of $ABS_{600}$ ≈ 0.8. ρ: Spearman correlation coefficient. Values for transcript levels are the average of duplicate independent experiments, while values for the depletion score are the average of duplicate independent screens.

(**B**) Predicted translation rate poorly correlates with guide depletion score in the library screen. Translation rates were predicted using the RBS calculator (Salis, 2011).

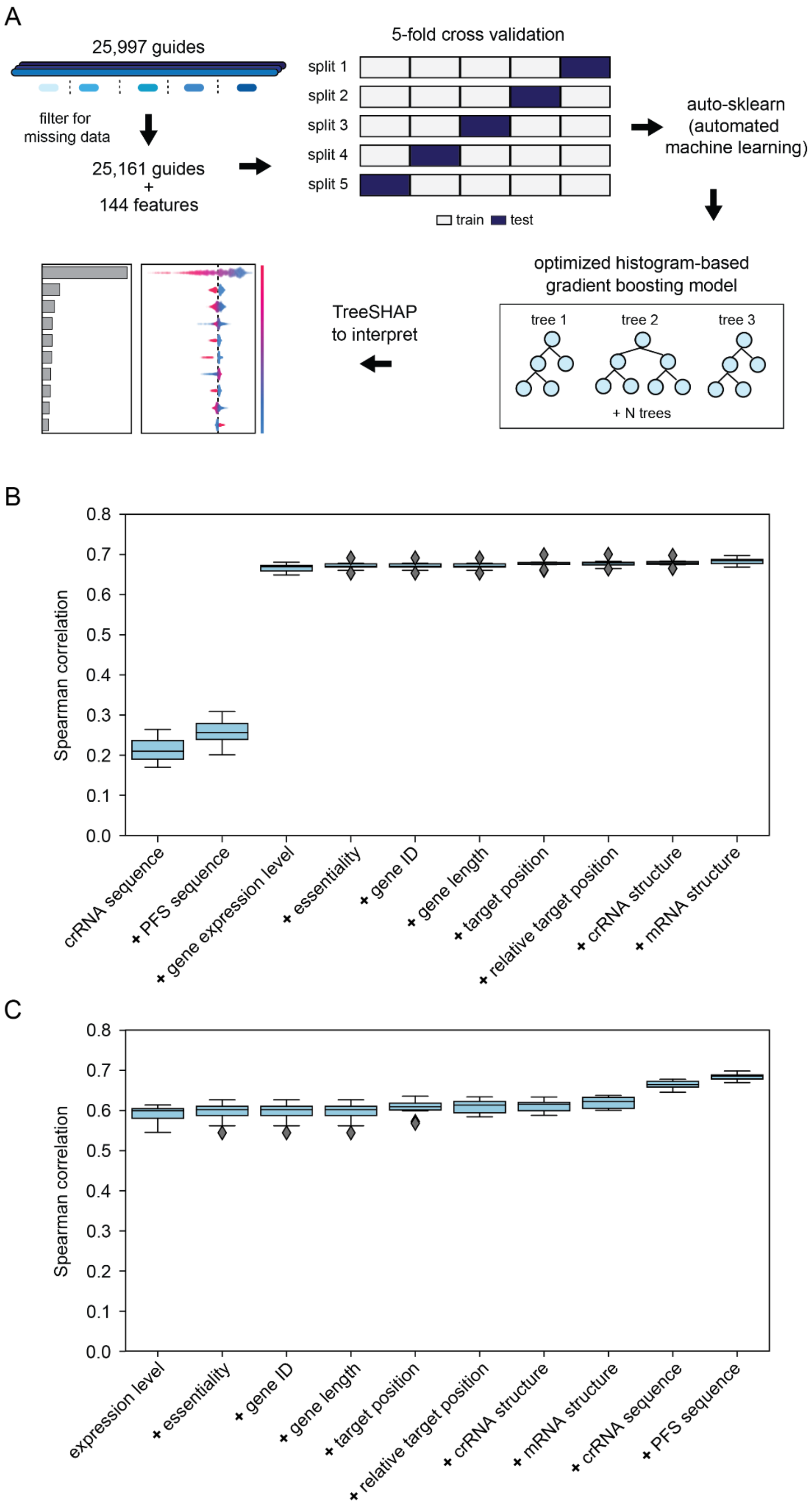


**Figure S6.** Machine learning model and feature importance. Related to **Fig. 3**.

(**A**) Overview of the machine learning model development protocol used. Screen data was used to determine the depletion of each of 25,161 guides as part of growth in liquid culture. The derived $\log_2$ fold-changes were used as prediction targets, with 144 sequence, gene, thermodynamic, and expression features as predictors. Auto-sklearn was used to train and optimize an assortment of candidate model types. A histogram-based gradient boosting model was the best performing model. This model was then interpreted using TreeSHAP to explain the contribution of each predictor to each prediction.

(**B**) Alternative evaluation of feature importance using cross-validation. The predictive power of each feature was evaluated by first training a simple histogram-based gradient boosting model with a single feature or feature set, and then iteratively adding additional features and comparing the Spearman correlation in 10-fold cross-validation of the model with the expanded feature set to previously trained models. For B, the base feature was the crRNA sequence. Adding PFS sequence features slightly increases the Spearman correlation, but a large jump in correlation is observed when adding expression level features. Other features lead to only minor changes in the Spearman correlation.

(**C**) As in B, but using expression level as the base feature. This again shows that expression level alone has high predictive utility; features other than the crRNA and PFS sequence features have relatively minor effects on overall prediction accuracy.

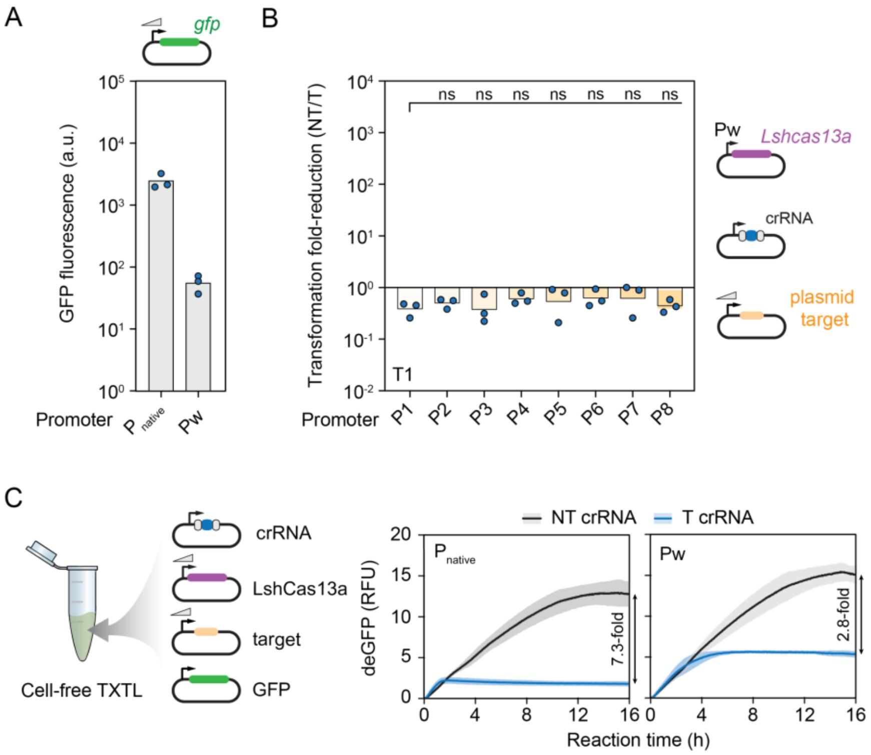


**Figure S7.** Lowering Cas13a expression levels eliminates immune activation even with high target expression.

(**A**) Quantified strength of the native promoter from *L. shahii* and the weaker constitutive Pw promoter driving expression of LshCas13a. Promoter strength was measured based on the fluorescence produced from a downstream *gfp* reporter.

(**B**) Impact of lowering Cas13a expression levels on immune activation in *E. coli*. Statistical significance was calculated by comparing the transformation fold-reduction to that of the weaker P1 promoter. ***: $p < 0.001$. **: $p < 0.01$. *: $p < 0.05$. ns: not significant.

(**C**) TXTL-based quantification of Cas13a targeting activity. The same constructs used in *E. coli* along with the non-targeted GFP reporter were added to a TXTL reaction, and collateral cleavage activity was measured over time based on reduction in GFP fluorescence.

Bars in A and B represent the mean of triplicate independent experiments. The lines and shaded regions in C represent the mean and standard deviation of triplicate experiments.

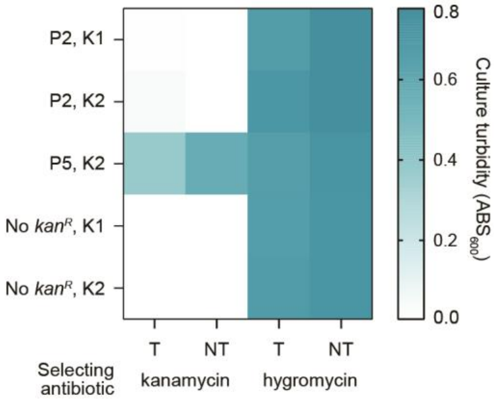


**Figure S8.** Heat map comparing growth in the presence of kanamycin (50 μg/mL) or hygromycin (100 μg/mL) for constructs associated with a low transformation fold-reduction in **Fig. 6B**. Related to **Fig. 6**.

Constructs lacking the *kan*$^R$ gene serve as negative controls, while the non-targeting crRNA plasmid serves as a non-targeting (NT) control. The same NT control was used for K1 and K2. Values represent turbidity measurements after 12 h of growth averaged across 9 biological replicates. One of the constructs allows growth on the higher kanamycin concentration despite active targeting by Cas13a.

### SUPPLEMENTARY TABLES

**Supplementary Table 1.** Anderson promoters used in this study. For other Anderson promoters, refer to http://parts.igem.org/Promoters/Catalog/Anderson**.** Transcription is predicted to begin after the 3′ end of the sequence.

| Promoter name | Promoter # | Sequence (5′ to 3′) |
|---|---|---|
| PJ23112 | P1 | CTGATAGCTAGCTCAGTCCTAGGGATTATGCTAGC |
| PJ23109 | P2 | TTTACAGCTAGCTCAGTCCTAGGGACTGTGCTAGC |
| PJ23117 | P3 | TTGACAGCTAGCTCAGTCCTAGGGATTGTGCTAGC |
| PJ23114 | P4 | TTTATGGCTAGCTCAGTCCTAGGTACAATGCTAGC |
| PJ23116 | P5 | TTGACAGCTAGCTCAGTCCTAGGGACTATGCTAGC |
| PJ23115 | P6 | TTTATAGCTAGCTCAGCCCTTGGTACAATGCTAGC |
| PJ23105 | P7 | TTTACGGCTAGCTCAGTCCTAGGTACTATGCTAGC |
| PJ23119 | P8 | TTGACAGCTAGCTCAGTCCTAGGTATAATGCTAGC |
| PJ23108 | Pw | CTGACAGCTAGCTCAGTCCTAGGTATAATGCTAGC |

**Supplementary Table 2.** Strains and plasmids used in this study. See SI_Tables.xlsx file, tab 1.

**Supplementary Table 3.** Oligos used in this study. See SI_Tables.xlsx file, tab 2.

**Supplementary Table 4.** Library design and oligos. See SI_Tables.xlsx file, tab 3.
